# Spatially Mapping the Mechanical and Structural Properties of the Seedling Uterine Fibroid-Myometrium Interface

**DOI:** 10.1101/2025.11.14.687752

**Authors:** Daniella M. Fodera, Alara Sutcu, Arielle S. Joasil, Aidan M. Therien, Johanna L. L. Jackson, Arnold P. Advincula, Xiaowei Chen, Christine P. Hendon, Michelle L. Oyen, Tal Korem, Kristin M. Myers

**Affiliations:** Department of Biomedical Engineering, Columbia University, New York, NY, USA; Institute of Human Nutrition, Columbia University Irving Medical Center, New York, NY, USA; Program for Mathematical Genomics, Department of Systems Biology, Columbia University Irving Medical Center, New York, NY, USA; Department of Electrical Engineering, Columbia University, New York, NY, USA; Department of Obstetrics & Gynecology, Columbia University Irving Medical Center, New York, NY, USA; Department of Pathology and Cell Biology, Columbia University Irving Medical Center, New York, NY, USA; Department of Biomedical Engineering, Wayne State University, Detroit, MI, USA; Department of Mechanical Engineering, Columbia University, New York, NY, USA

**Keywords:** uterine fibroids, leiomyomas, myometrium, biomechanics, tissue interface, microindentation, time-dependence, permeability

## Abstract

Uterine fibroids (leiomyomas) are highly prevalent, noncancerous tumors canonically described as stiff and collagen-dense. Still, the mechanical heterogeneity within uterine fibroids and the alterations that occur at the fibroid–myometrium interface remain poorly understood. This study quantitatively maps the local mechanical, structural, and compositional properties of seedling uterine fibroids (*<* 1 cm) at the interface region. Relative to patient-matched myometrial tissues, uterine fibroids exhibited increased stiffness, decreased permeability, increased diffusivity, decreased hydration, greater collagen content, and distinct collagen organization. At the fibroid–myometrium interface, a band of aligned myometrial fibers immediately adjacent to the fibroid was observed, and heterogeneous spatial patterns in elastic modulus and permeability were identified. Ultimately, this study establishes foundational knowledge on the mechanics and structure of seedling uterine fibroids, facilitating future developments of clinically translatable detection tools.

## Introduction

Uterine fibroids, also known as leiomyomas, are noncancerous tumors that form within the uterus and are present in up to 70–80% of women by age 50, affecting an estimated 26 million individuals in the United States^1–4^. Among those affected, 10–50% are reported to experience clinically significant symptoms in their reproductive years^5–7^. Uterine fibroids may cause prolonged heavy menstrual bleeding, chronic pelvic pain, anemia, and infertility, as well as pregnancy complications in the form of miscarriage, severe acute pelvic pain, preterm birth, and placental abnormalities^4,6,8,9^. Fibroids are known to be highly responsive to hormones, specifically estrogen^10–12^. Fibroids begin forming after menarche (puberty) and peak in prevalence and symptomology during the peri-menopausal years, notably decreasing with parity (number of pregnancies) and in menopause^11,13^. It has been hypothesized that normal reproductive events such as ovulation, menstruation, and embryo implantation may be primary initiators for this disease, leading to genetic mutations of smooth muscle cells and thereby driving fibroid formation^12,14,15^. Still, despite their prevalence, the etiology and pathophysiology of uterine fibroids remain poorly understood^15^. As a result, the disease burden is high from both a quality of life and an economic perspective; direct and indirect costs of this disease in the United States have been estimated between $5.9–34.4 billion annually^16^.

Treatment options for uterine fibroids are limited and lag in comparison to other nonmalignant diseases, due in part to historic underfunding^15,17,18^. Surgery has long been the most common and effective method for uterine fibroid treatment, but the only definitive method to eliminate and prevent the re-growth of uterine fibroids is a total hysterectomy – the surgical removal of the entire uterus^6,15,19,19^. For individuals seeking to preserve their fertility, myomectomies and uterine artery embolizations are two procedures clinicians regularly employ to surgically excise or halt blood flow to uterine fibroids, respectively^6,15,19^. While these procedures can alleviate the severity of an individual’s symptoms, the benefit is often temporary, and the recurrence rates of uterine fibroids are estimated to be between 12–60% within one to five years post-surgery^20–24^. Failure to completely remove all uterine fibroids can contribute to disease recurrence^22^. Seedling fibroids (*<* 1 cm) are particularly difficult to detect and remove intraoperatively due to their small size and often large quantity^25^. Pre-operative imaging modalities (e.g., ultrasound, MRI) lack the fine resolution needed to identify seedling fibroids^25,26^, and surgical technique largely relies on manual palpation and gross visual assessment to identify stiff, bright white, or abnormally shaped masses^27,28^.

Uterine fibroids are known to exhibit immense heterogeneity, varying widely in size, location, and gross architecture within and across different patients. Though they can regularly grow beyond 5 cm in diameter^29,30^, occasionally reaching 50 cm^31–33^, all originate as seedling fibroids on the millimeter scale^6,34^. Seedling uterine fibroids represent a crucial stage in fibroid development, offering insight into the complex growth and remodeling patterns of these tumors. Fibroids can be broadly classified by their location of growth (Fig. 1a)^6,34–36^. Intramural (IM) fibroids are the most common type, present within the myometrium; submucosal (SM) fibroids form adjacent to the endometrium and subserosal (SS) fibroids adjacent to the perimetrium (serosa), protruding into the intrauterine or abdominal cavity, respectively (Fig. 1a)^35–39^. In addition, rare variants of uterine fibroids can present as hard, calcified masses or soft, spongy, degenerating tissues, often due to age or changes in their vascular supply^40–44^.

**Figure 1.**
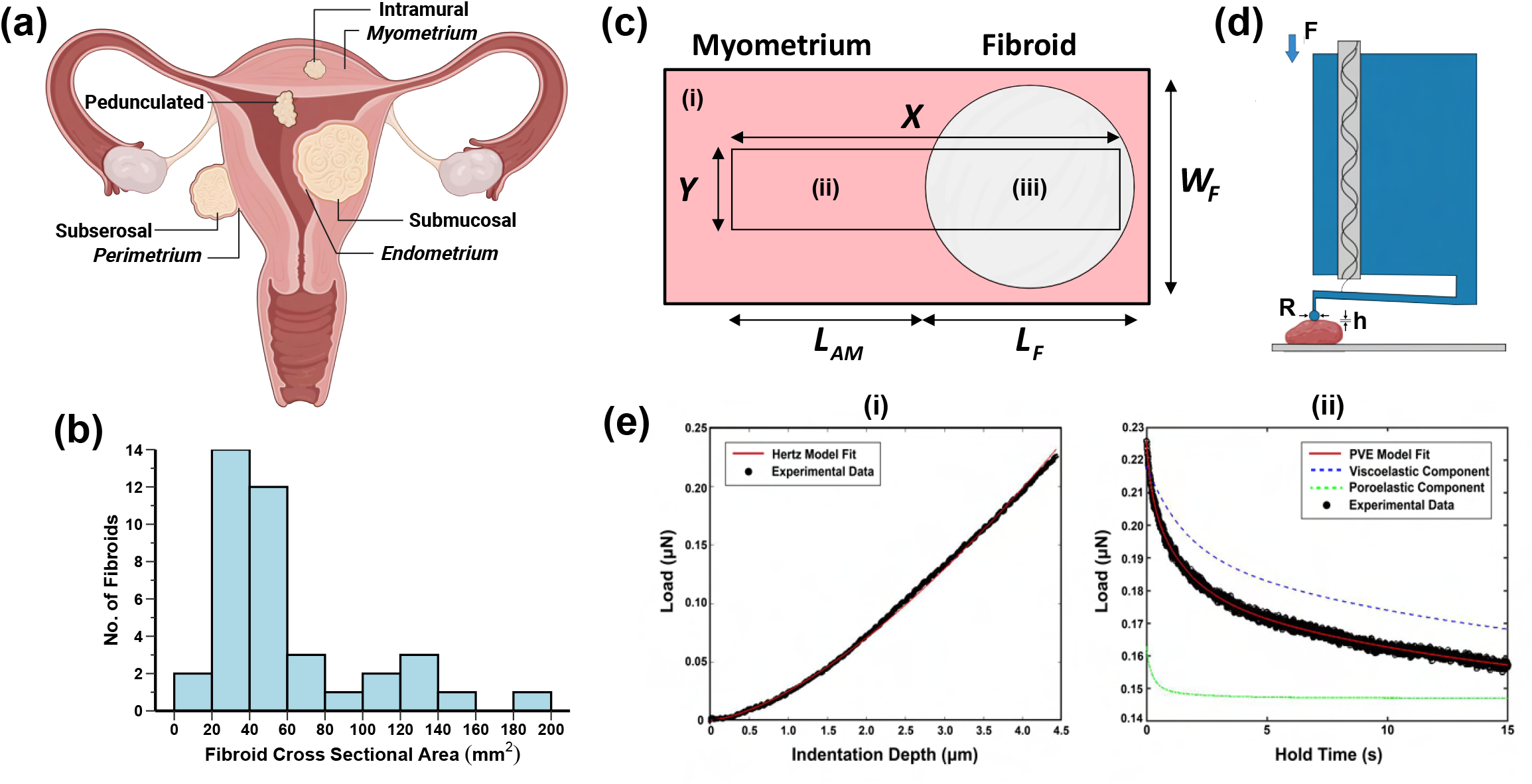
Summary of fibroid sample characteristics and microindentation testing approach. **(a)** Schematic of the female reproductive tract illustrating uterine tissue layers (i.e., endometrium, myometrium, and perimetrium) and fibroid subtypes (i.e., submucosal, intramural, subserosal). Created with Biorender.com. **(b)** Distribution of seedling uterine fibroid sizes (i.e., cross-sectional areas) analyzed in this dataset. **(c)** Approach for spatially mapping the fibroid-myometrium interface. Three tissue types were examined: (i) distant myometrium, (ii) adjacent myometrium, and (iii) fibroid. Fibroid length (*L*_*F*_), fibroid width (*W*_*F*_), and adjacent myometrium length (*L*_*AM*_) are labeled. X and Y indicate the entire length and width of the rectangular testing region, respectively. **(d)** Schematic of the Piuma nanoindentor (Adapted from Optics11 Life). A spherical probe with a radius (*R*) of 50 *µ*m is attached to the end of a cantilever and is indented into the sample to a fixed depth (*h*) of 4 *µ*m, recording load (*F*) over time. **(e)** Representative data generated from microindentation testing were fit with the (i) Hertzian contact model and (ii) a combined poro-viscoelastic (PVE) model. Data are shown for the (i) loading and (ii) hold portions of the testing protocol.

Structurally and compositionally, uterine fibroids differ markedly from myometrial tissue and exhibit size-dependent variations. The myometrium is the thickest layer of the uterus and is primarily composed of smooth muscle fascicles interspersed with collagen and elastin fibers^38,39,45^. Fibroids share these core cellular and extracellular matrix (ECM) constituents, but differ from the myometrium in their relative proportions^34,46^. Interestingly, the cellular makeup of uterine fibroids appears to be size dependent; smaller fibroids have been found to be more cellular, with higher SMC-to-fibroblast ratios, than larger tumors^46^. Abundant ECM deposition, driven largely by increased collagen content, is a defining hallmark of fibroids^34,47–49^. Still, specific ECM components reported to differ between fibroid and myometrium tissues (i.e., collagen sub-types, glycosaminoglycans, and proteoglycans) vary widely in the literature^34,47–50^. Beyond compositional differences, fibroids exhibit altered ECM organization^34^. Uterine fibroid architecture has been classically described as whorled, but this characterization scheme has recently been expanded to describe greater fibroid heterogeneity^34^. Notably, Jayes et al. (2019) identified at least four distinct tissue patterns (i.e., whorled, nodular, interweaving trabecular, and mixed) for fibroids at the centimeter length scale based on qualitative observations of tissue architecture^34^. The manner in which uterine fibroids affect the composition and structure of the normal surrounding myometrium is not deeply understood and is a matter of debate.

Increased biomechanical stiffness is another distinguishing feature of uterine fibroids^34,47,48,51,52^. At a fundamental level, the mechanical properties of tissues are governed by their underlying structure and composition, and alterations to these features can drive abnormal cell behavior via mechanotransduction, further contributing to disease progression^53–56^. Mechanically, uterine fibroids have been found to be 1.5–5 times stiffer than patient-matched myometrial tissues^34,47,48,51,52,57–59^. Both uterine fibroid and myometrium tissues have also been shown to exhibit time-dependent material behavior^45,57,59–62^, with distinctions in tissue types observed under compression and shear^57,59,60^. Still, the relative contribution of viscoelastic (rearrangement of the solid matrix) and poroelastic (pressurization and motion of fluid through pores) has yet to be considered in the context of fibroids^63–65^. Further, *ex vivo* mechanical tests have largely characterized the bulk properties of centimeter-sized uterine fibroids under compression and shear, resulting in a single measurement per sample^34,47,51,57,59^. As such, the mechanical properties of seedling uterine fibroids and their local heterogeneity at the fibroid–myometrium interface remain uncharacterized.

This study quantifies the mechanical properties of seedling uterine fibroids, characterizing both intra- and inter-sample heterogeneity at the micro-to milli-meter length scales. Specifically, this study spatially maps the distribution of mechanical, structural, and compositional properties across the fibroid-myometrium interface for seedling uterine fibroids ≤ 1 cm in diameter using microindentation, histology, and optical imaging approaches. Ultimately, this comprehensive spatial characterization of the structure-function relationship of seedling uterine fibroids will (i) deepen fundamental understanding of fibroid pathophysiology, including growth and remodeling mechanisms, (ii) aid in the development of novel intraoperative margin detection tools, and (iii) inform therapeutic delivery approaches for pharmacological intervention.

## Results

### Micro-Mechanical Characterization of the Uterine Fibroid-Myometrium Interface

Thirty-eight seedling uterine fibroids, all less than 1 cm in diameter and embedded in myometrial tissue, were obtained from total hysterectomy procedures from twelve nonpregnant human participants, aged 36–49 (Tables 1, 2). Detailed patient information is noted in Table S1 and Fig. S1. Five patients were classified as nulliparous (parity = 0) and seven as parous (parity ≥ 1). This dataset includes 6 submucosal (SM), 32 intramural (IM), and 1 subserosal (SS) fibroids sampled from three anatomic regions of the uterus (i.e., anterior, fundus, posterior); all fibroid samples were validated histologically by a clinical pathologist (Tables 2, S2; Fig. S2). Fibroid size, calculated as the area of an ellipse, was on average 49.5 *mm*^2^ with a standard deviation of 41.3 *mm*^2^ (Fig. 1b).

**Table 1.** Summary of patient demographics.

| Characteristic | N = 12 | (%) |
| --- | --- | --- |
| Age (years) | Mean ± SD: $44 \pm 4$<br>Range: 36–49 | |
| Parity | Nulliparous: 5<br>Parous: 7 | (41.7)<br>(58.3) |
| Gravidity | Median: 1.5<br>Range: 0–3 |  |
| Estimated Menstrual Cycle Stage | Proliferative: 4<br>Secretory: 2<br>Inactive: 2<br>Unknown: 3<br>N/A. Post-menopausal: 1 | (33.3)<br>(16.7)<br>(16.7)<br>(25.0)<br>(8.3) |
| Hormone Therapy | Yes: 3<br>No: 9 | (25.0)<br>(75.0) |

*Number of subjects, N, and percentage of total subjects (in parentheses) in each group. Gravidity is defined as the total number of pregnancies, regardless of length, and parity is the total number of pregnancies that have progressed past 20 weeks of gestation resulting in a live birth.*

**Table 2.** Summary of characteristics for seedling fibroids evaluated in this dataset.

| Category | N = 38 specimens | (%) |
| --- | --- | --- |
| **Specimens per patient** | Median (range): 2 (1–8) |  |
| **Anatomic region** | Anterior: 19 | (50.0) |
|  | Posterior: 10 | (26.3) |
|  | Fundus: 6 | (15.8) |
|  | Unknown: 3 | (7.9) |
| **Fibroid subtype** | Intramural: 32 | (84.2) |
|  | Submucosal: 5 | (13.2) |
|  | Subserosal: 1 | (2.6) |

*Number of specimens (N) and percentage of total specimens indicated for each group.*

To investigate whether uterine fibroids alter the material properties of the immediate surrounding myometrium, two groups of myometrial tissues were assessed in this study: (i) adjacent myometrium (≤ 3 mm from at least one visible fibroid) and (ii) distant myometrium (*>* 5 mm from any visible fibroid) (Fig. 1c). Spherical microindentation (*R* = 50 *µ*m, *h* = 4 *µ*m) was employed to measure the time-dependent (i.e., viscoelastic and poroelastic) material properties of uterine fibroids and patient-matched myometrial tissues within the linear elastic regime (Fig. 1d). Each sample containing uterine fibroid and adjacent myometrium tissue was tested as a rectangular region, shown in Fig. 1c, and was scaled according to the individual fibroid’s dimensions. The length (*X*) of the testing region was equivalent to the entire length of the fibroid (*L*_*F*_) plus a fixed 3 mm of the adjacent myometrium (*L*_*AM*_); the width was equal to 40% of the fibroid’s width (*W*_*F*_). Measurements were taken every 200 *µ*m in the *x* and *y* directions along the tissue’s surface. For each patient, three samples of distant myometrial tissue, taken from three anatomic regions, were characterized with approximately 150 discrete, non-overlapping indentation points to capture intra-sample heterogeneity. All in all, this dataset represents more than 12,000 individual indentation measurements across all three tissue types. To assess the intrinsic elasticity, viscoelasticity, and poroelasticity of the uterus, these data were fit to a Hertzian contact model^66^ and a combined poroelastic-viscoelastic (PVE) model^67^ to determine the following parameters: apparent elastic modulus (*E*_*r*_), instantaneous elastic modulus (*E*_0_), equilibrium elastic modulus (*E*_∞_), poroelastic modulus (*E*_*PE*_), viscoelastic ratio (*E*_∞_*/E*_0_), intrinsic permeability (*k*), and diffusivity (*D*). Representative mechanical data analyzed by these two models are shown in Fig. 1e. For specimens containing paired fibroid and adjacent myometrium data, the relative change in material parameters was assessed for (i) all data points (Δ[*E*_*r*_]_*all*_, Δ[*E*_∞_*/E*_0_]_*all*_, Δ*k*_*all*_, Δ*D*_*all*_) and (ii) those immediately surrounding (± 200 *µm*) the interface region (Δ*E*_*int*_, Δ*k*_*int*_).

### Material Properties by Tissue Type

For all elastic modulus measures evaluated (*E*_*r*_, *E*_0_, *E*_∞_, *E*_*PE*_), uterine fibroids were stiffer (*E*_*r*_ = 2.87 ± 1.90 kPa) than both adjacent (*E*_*r*_ = 1.00 ± 0.40 kPa) and distant myometrium (*E*_*r*_ = 1.05 ± 0.46 kPa) at the scale of microindentation testing (Figs. 2a, S3). Statistical significance was determined using a linear mixed model, which was applied to all subsequent analyses unless otherwise noted, with p-values reported in the corresponding figures. There were no significant differences in the elastic modulus measures of adjacent and distant myometrial tissues (Fig. 2a). Relative to myometrial tissues, fibroids were stiffer by 3.47 ± 3.6 fold (Fig. 2b). Each elastic modulus parameter differed slightly in its absolute value but retained the same overall trends across tissue types (Fig. S3).

**Figure 2.**
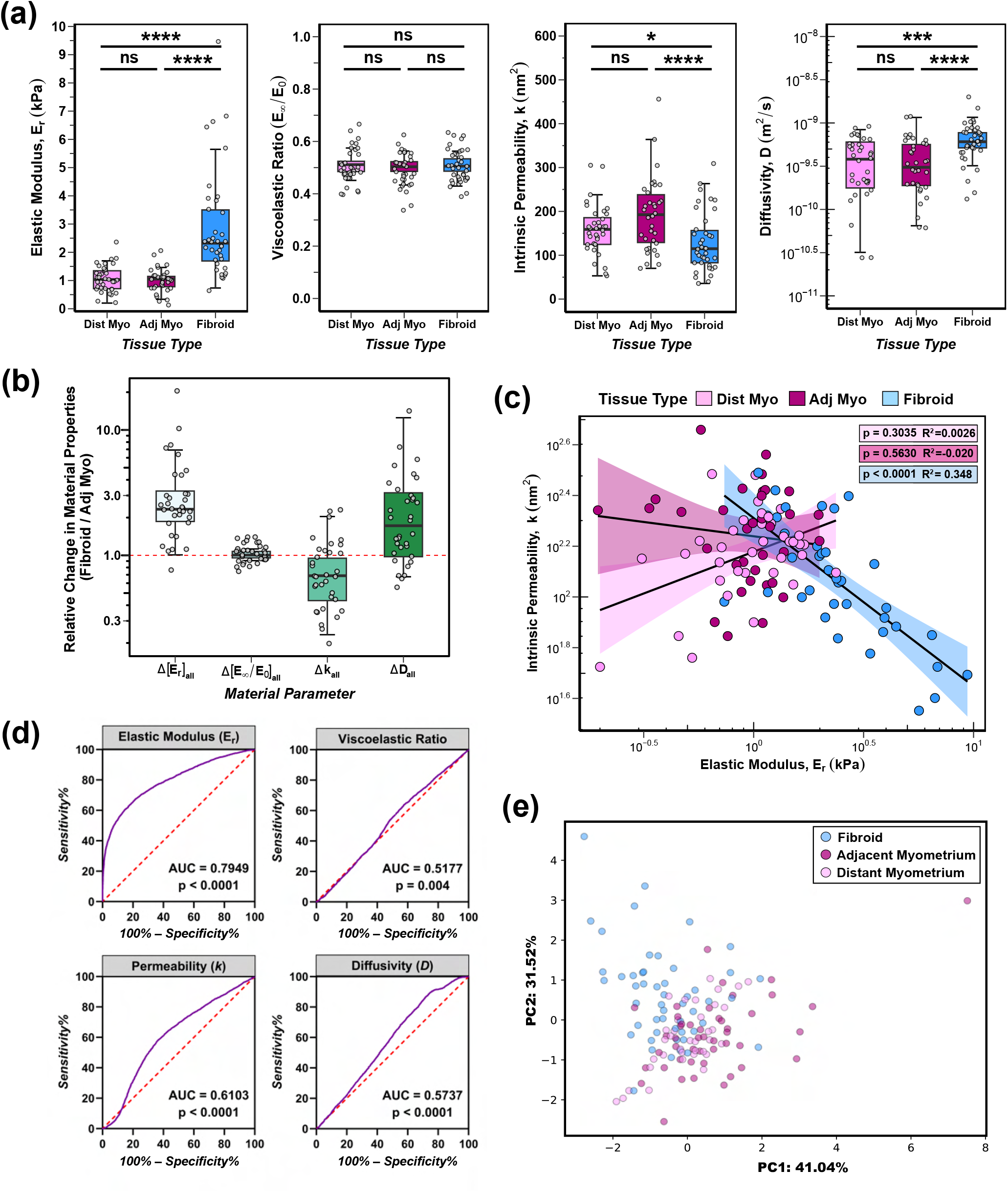
Seedling uterine fibroids exhibit distinct mechanical signatures from patient-matched myometrial tissues. **(a)** Material properties by tissue type: elastic modulus (*E*_*r*_), viscoelastic ratio (*E*_∞_*/E*_0_), intrinsic permeability (*k*), and diffusivity (*D*). Each symbol represents the median value of all indentation points for a given sample. Statistical analysis was performed with a linear mixed model with significance marked as follows: ^ns^ *p >* 0.05, ^***^ *p <* 0.001, ^****^ *p <* 0.0001. **(b)** Relative fold change in material properties for fibroid tissues relative to matched adjacent myometrial tissue. A red, horizontal dashed line at 1.0 is included for reference to indicate no fold change. **(c)** Intrinsic permeability (*k*) versus elastic modulus (*E*_*r*_) plotted on a log-log scale. Each symbol represents the median value of all indentation points measured for an individual sample. Linear regression analysis was performed for each tissue type; shaded regions indicate the 95% confidence interval, with corresponding *p* values shown. **(d)** Receiver operating characteristic (ROC) curves (*purple*), plotting sensitivity versus specificity, for all material properties (*E*_*r*_, *E*_∞_*/E*_0_, *k, D*) were used to predict differences between fibroid and myometrium (adjacent & distant) tissues. Area under the curve (AUC) and *p* values from null-hypothesis testing are noted for each comparison. The closer an AUC value is to 1, the better the classifier is^68^. A dashed red line is included for reference to indicate random chance with no predictive value (AUC = 0.5). **(e)** Principal component analysis (PCA) considering all material properties for a given sample. Each symbol represents a distinct tissue specimen; data is colored by tissue type.

Mean viscoelastic ratio (*E*_∞_/*E*_0_) across all tissue types was equivalent to 0.50 ± 0.11, therefore indicating the mechanical behavior of uterine tissue is time-dependent (Fig. 2a). There was a small, statistically significant difference in viscoelastic ratio (*E*_∞_/*E*_0_) between adjacent myometrium (0.496 ± 0.043) and fibroid (0.508 ± 0.035) tissue types; no other statistical significance across tissue types was noted for this parameter (Fig. 2a). It is important to note, however, that the relative change in the viscoelastic ratio between fibroid and adjacent myometrial tissues was close to 1 (Figs. 2a,b).

Intrinsic permeability (*k*) and diffusivity (*D*) notably varied across tissue types (Fig. 2a). Fibroids exhibited decreased permeability (130.0 ± 67.5 *nm*^2^) compared to adjacent (192.3 ± 84.2 *nm*^2^) and distant (158.5 ± 56.6 *nm*^2^) myometrium tissues by 0.78 ± 0.46 fold (Fig. 2a,b). Pore size (*ξ*), derived from values of permeability 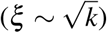, was found to be on the order of 10^1^ nm (Fig. S5). Additionally, fibroids exhibited increased diffusivity (6.86 ± 4.1 x 10^−10^ *m*^2^*/s*) compared to adjacent (3.99 ± 2.7 x 10^−10^ *m*^2^*/s*) and distant (3.91 ± 2.4 x 10^−10^ *m*^2^*/s*) myometrium tissues by 2.62 ± 2.44 fold (Fig. 2a,b). Inter-sample and intra-sample heterogeneity was identified for the mechanical properties of fibroids evaluated in this dataset. Among the 34 fibroids with matched adjacent myometrial tissue, 32 (94.1%) fibroids exhibited an increase in elastic modulus (Δ[*E*_*r*_]_*all*_ *>* 1), 29 (85.3%) exhibited no change in viscoelastic ratio (Δ[*E*_∞_*/E*_0_]_*all*_ ≃ 1), 24 (70.5%) exhibited a decrease in permeability (Δ*k*_*all*_ *<* 1), and 23 (67.6%) exhibited an increase in diffusivity (Δ*D*_*all*_ *>* 1) relative to myometrium tissue (Fig. S6). Intra-sample heterogeneity was quantified by computing the coefficient of variation (CV) across mechanical replicates for each sample. Fibroids displayed a markedly greater spread in elastic modulus values compared to both adjacent and distant myometrium tissues, suggesting a higher degree of mechanical heterogeneity (Figs. S3, S4a). Elastic modulus values ranged from approximately 50 Pa to 80 kPa for fibroids and 20 Pa to 30 kPa for myometrium specimens. Conversely, fibroids demonstrated lower variability in viscoelastic ratio, as reflected by reduced CVs, and displayed similar degrees of variability for permeability and diffusivity metrics across all tissue types (Fig. S4).

The predictive value of individual material parameters (*E*_*r*_, *E*_∞_/*E*_0_, *k, D*) to distinguish between distinct tissue types was then assessed with receiver operating characteristic (ROC) curves (Fig. 2d). Elastic modulus was shown to be a fairly good predictive measure in classifying fibroid versus myometrial tissues, with a corresponding AUC value of 0.79 (Fig. 2d). Viscoelastic ratio, permeability, and diffusivity were alone poor predictive measures to differentiate between fibroid and myometrial tissues, as evidenced by AUC values of 0.52, 0.61, and 0.57, respectively (Fig. 2d). All AUC values computed between adjacent and distant myometrium groups were less than 0.57, indicating these tissues are poorly differentiated from one another on an individual material property basis (Fig. S8). Principal component analysis (PCA) considering all measured material properties revealed clear separation between fibroid and myometrium tissues (Fig. 2e). The PCA points of distant and adjacent myometrial tissues largely overlapped (Fig. 2e).

### Correlation of Distinct Material Parameters

Uterine fibroids exhibited a negative linear relationship between permeability and elastic modulus on a log-log scale (Figs. 2c, S9). Neither adjacent nor distant myometrium tissues exhibited any statistically significant correlation between elastic modulus and permeability values (Fig. 2c). For all tissue types evaluated, there were no correlations found between (i) viscoelastic ratio and elastic modulus and (ii) viscoelastic ratio and permeability (Fig. S10). Of note, diffusivity was not evaluated in this manner due to its mathematical dependence on elastic modulus and permeability values.

### Effect of Tissue Characteristics on Micro-Mechanical Properties

Comparing across fibroid subtypes, there were no differences in any material properties evaluated between submucosal and intramural fibroids; no comment can be made for subserosal fibroids due to the small sample size (n = 1) of this group (Fig. 3a). Investigating correlations between fibroid size and mechanical properties, there was a small, but statistically significant (*p* = 0.0036, *R*^2^ = 0.195) negative correlation between fibroid size and viscoelastic ratio (Fig. 3b). No other material parameters (i.e., elastic modulus, permeability, and diffusivity) were correlated with fibroid size (Fig. 3b). Additionally, for the three tissue types tested, there were no systemic differences in material properties across the anterior, fundus, and posterior regions sampled (Fig. S11).

**Figure 3.**
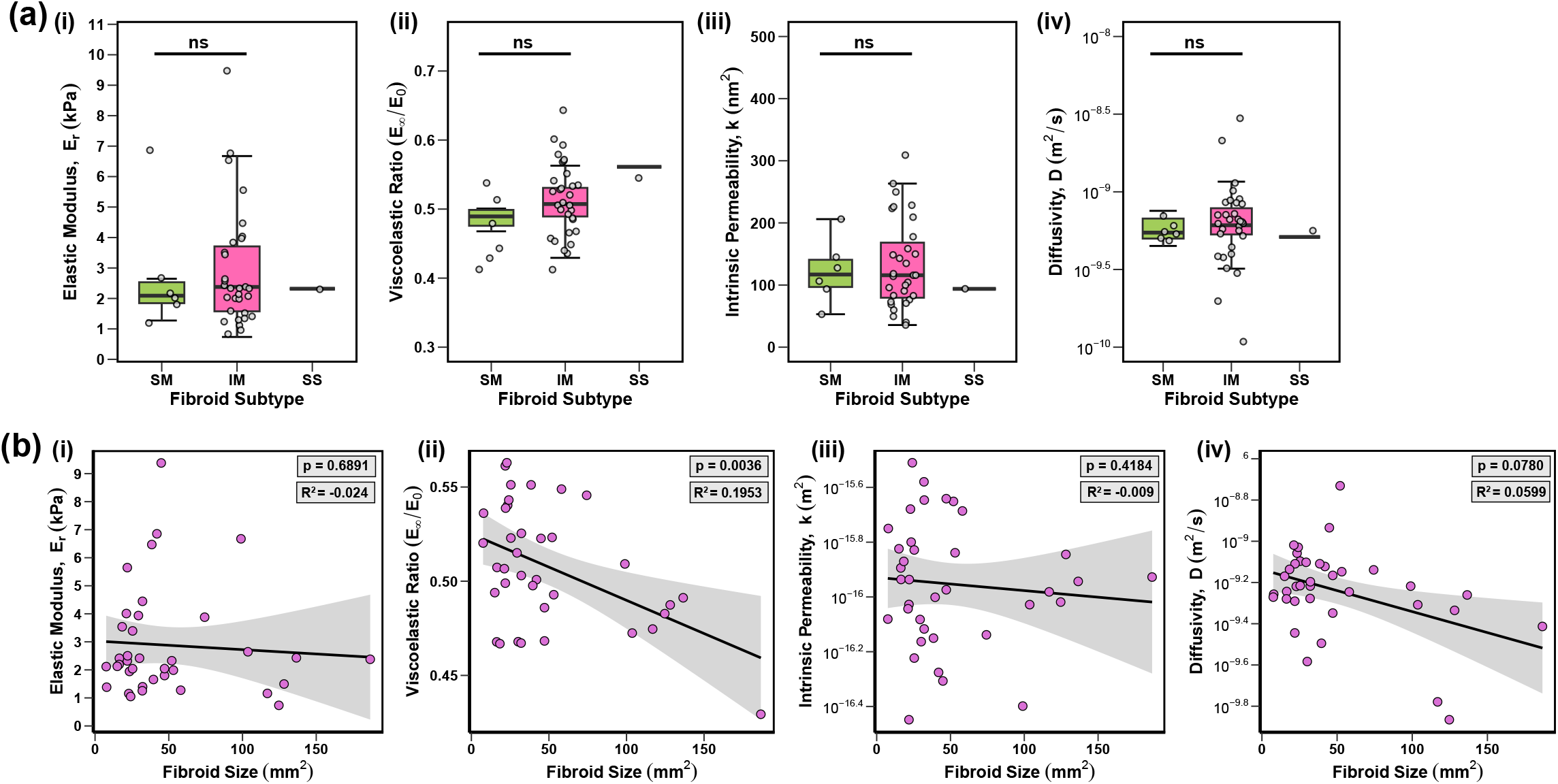
Fibroid size and subtype were variably associated with the material properties of seedling uterine fibroids. **(a)** Variations in (i) elastic modulus (*E*_*r*_), (ii) viscoelastic ratio (*E*_∞_*/E*_0_), (iii) intrinsic permeability (*k*), and (iv) diffusivity (*D*) values with fibroid subtype (i.e., submucosal [SM], intramural [IM], subserosal [SS]). Statistical analysis was performed with a linear mixed model, where ^ns^p *>* 0.05. **(b)** Correlations between fibroid size (i.e., cross-sectional area) and material properties (*E*_*r*_, *E*_∞_*/E*_0_, *k, D*). Each symbol represents the mean value of all indentation points measured for an individual sample. The p-values and adjusted *R*^2^ values from linear regression analyses are noted in the upper right corner of each plot. For (a) and (b), each symbol represents the median value of all indentation points measured for an individual sample.

### Effect of Patient Characteristics on Micro-Mechanical Properties

All three tissue types exhibited some degree of patient-specific variability for all material parameters (Fig. S12). Statistically significant changes as a function of parity and age were found for a subset of material properties and tissue types (Figs. 4, S13). Parity and age had no association with the elastic moduli of all three tissue types, but variable trends were shown for viscoelastic ratio, permeability, and diffusivity (Figs. 4, S13). With regard to viscoelastic ratio, distant myometrium samples derived from parous patients exhibited a slight decrease in viscoelastic ratio relative to nulliparous counterparts; this trend is absent for adjacent myometrium and fibroid tissue types (Fig. S13). Further, the viscoelastic ratio for distant myometrium exhibited a slight positive correlation with age (Fig. 4). Intrinsic permeability was decreased for fibroids taken from parous patients relative to nulliparous individuals; no differences were seen for both myometrium groups (Fig. S13). Notably, there was a statistically significant positive correlation between age and permeability for the distant myometrium, but this same trend was not found for the adjacent myometrium (Fig. 4). Lastly, diffusivity exhibited a statistically significant positive correlation with age for fibroids, but not for adjacent or distant myometriums (Fig. 4). Diffusivity did not change with parity across all three tissue types (Fig. S13). Of note, statistical analyses between material properties and patient characteristics such as menstrual cycle stage, menopausal status, race, and ethnicity were not formally evaluated for this dataset due to small sample sizes (Figs. S1, S14).

**Figure 4.**
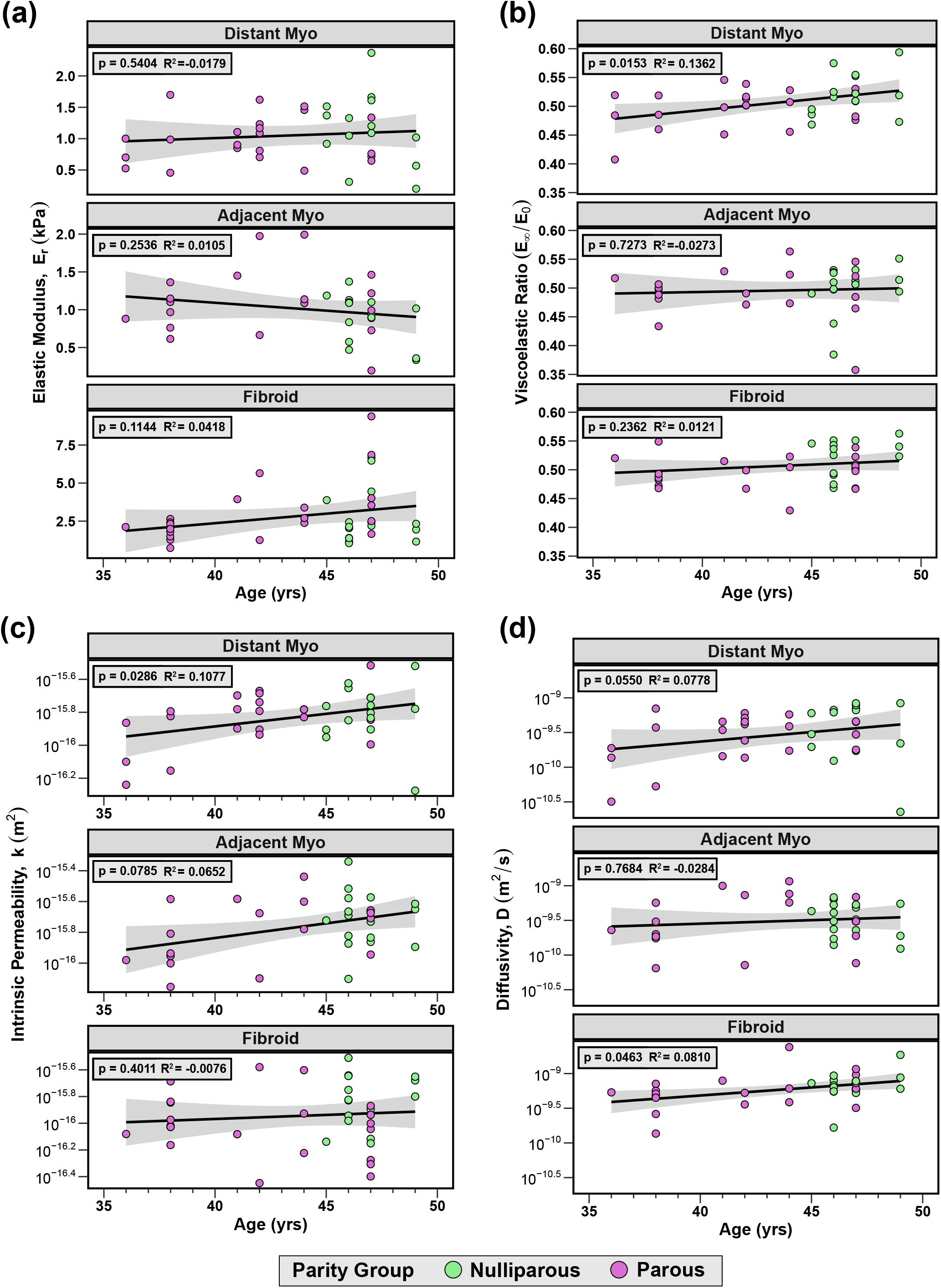
Age and parity show variable correlations with the material properties of distant myometrium, adjacent myometrium, and fibroid tissues. Each symbol represents the mean value of all indentation points measured for an individual sample. Colored circles indicate parity status: nulliparous (green) and parous (pink). The p-values and adjusted *R*^2^ values corresponding to each linear regression analysis are noted in the upper left corner of each plot. Data for elastic modulus and viscoelastic ratio are plotted on a linear scale; permeability and diffusivity on a logarithmic scale.

### Spatial Assessment of Material Properties Across the Uterine Fibroid-Myometrium Interface

Peak values of elastic moduli across the fibroid-myometrium testing region were largely exhibited by fibroid tissues (Fig. S15). For each individual fibroid, the locations (Δ*x*_*norm*_, Δ*y*_*norm*_) of the maximum and minimum elastic modulus (*E*_*r*_) values, normalized to the dimensions of the tested fibroid region, were identified (Fig. 5b). Extrema measurements were found across the entire width and length of the fibroid testing region and were not localized to the center of the fibroid (Figs. 5b, S16). Similar analysis revealed that peak permeability values occurred across the entire length of both adjacent myometrium and fibroid tissues (Fig. S17).

**Figure 5.**
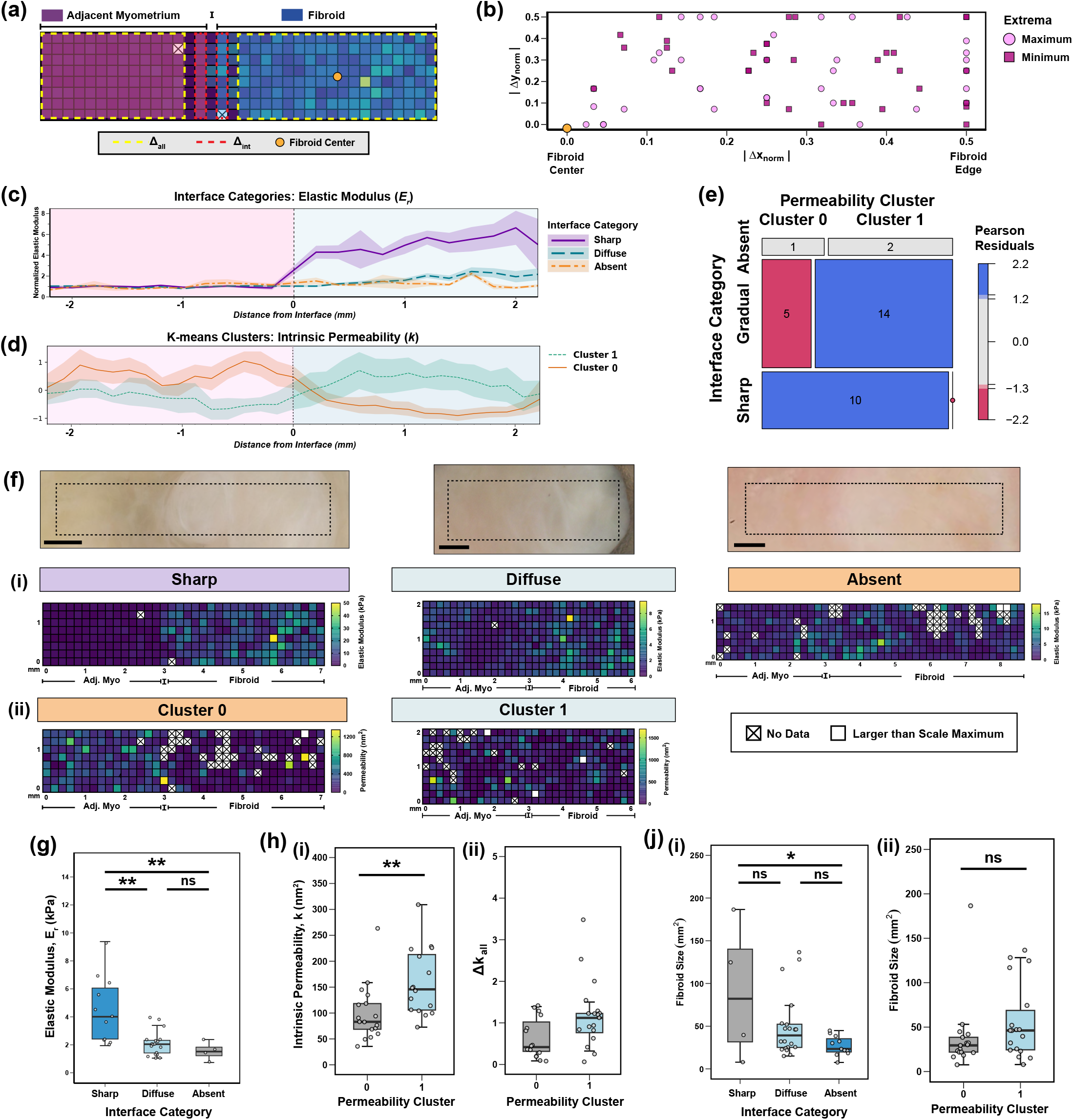
Distinct spatial mechanical gradients identified across the seedling uterine fibroid–myometrium interface. **(a)** Schematic illustrating the calculation of Δ_*all*_ and Δ_*int*_ parameters for all material properties. Adjacent myometrium, interface (I), and fibroid regions are labeled. An orange circle marks the fibroid’s center. Data included in the final calculations of Δ_*all*_ and Δ_*int*_ are shown by the yellow and red dashed lines, respectively. **(b)** Locations of minimum (square) and maximum (circle) elastic modulus (*E*_*r*_) values, determined relative to the fibroid’s center, are plotted for each sample. Data were normalized relative to the fibroid’s length (*L*_*F*_) and testing region width (*Y*). **(c, d)** Summary plots of identified mechanical interface categories. **(c)** Three interface categories have been identified for elastic modulus: sharp, gradual, and absent. Plotted data have been normalized by the median value of the adjacent myometrium region. Shaded regions represent standard error (s.e.m) values. **(d)** Two interface categories have been identified for permeability values based on k-means clustering. Shaded regions indicate the 95% confidence interval. Arbitrary units are shown on the y-axis. For panels **(c)** and **(d)**, the adjacent myometrium regions are shaded in pink and include negative X values. Fibroid regions are shaded in blue and include positive X values. A vertical dashed line at x = 0 marks the position of the interface. **(e)** Association between elastic modulus and permeability interface categories illustrated with a mosaic plot. Box area reflects the proportion of samples within each category pair, and color indicates Pearson residuals from the chi-square test of independence. Blue shading denotes combinations that occurred more frequently than expected under independence, whereas red indicates fewer than expected. All sharp interfaces overlapped with permeability cluster 0; gradual and absent interfaces predominantly overlapped with permeability cluster 1. **(f)** Representative samples exhibiting distinct interface patterns for (i) elastic modulus and (ii) permeability. Representative data corresponds to the following specimens: sharp & cluster 0 (NP1.F8), gradual & cluster 1 (NP10.F6.1), absent (NP2.F1). For each fibroid, a brightfield microscopy image of the fibroid-myometrium tested region; a scale bar representing a 1 mm distance is denoted by a solid black line in the lower left corner and the approximated region of fibroid-myometrium mapping is marked with a dashed black line. Matched spatial mechanical data for elastic modulus (*E*_*r*_) and permeability (*k*) is shown for each category; each square represents a distinct indentation point spaced 200 *µ*m apart. **(g)** Variations in elastic modulus values across identified interface categories (sharp, gradual, and absent). **(h)** Variations in the absolute (*k*) and relative (Δ*k*_*all*_) permeability values across permeability clusters. **(j)** Relationship between fibroid size and interface categories for (i) elastic modulus and (ii) permeability. Statistical analysis for panels (g), (h), and (j) was performed with a linear mixed model with significance marked as follows: ^ns^ *p >* 0.05, ^*^ *p <* 0.05), ^**^ *p <* 0.01). Each symbol represents a distinct fibroid sample.

At the interface region, on average, fibroids exhibited a 2.08 ± 1.93 and 0.93 ± 0.71 fold change in elastic modulus (Δ*E*_*int*_) and permeability (Δ*k*_*int*_) measures, respectively, relative to adjacent myometrium tissue (Fig. S18). For paired samples, the relative change in elastic modulus between fibroid and adjacent myometrium tissues varied depending on the location of analysis; (Δ*E*_*all*_) was larger than (Δ*E*_*int*_) and the relationship between these two parameters was not perfectly linear (Fig. S18). In contrast, for permeability measurements, there was no statistically significant difference between Δ*k*_*all*_ and Δ*k*_*int*_, and the relationship between these two parameters was linear (Fig. S18).

Given the notable spatial heterogeneity in elastic modulus values with localized changes at the interface, three distinct spatial patterns describing the transition in tissue stiffness at the fibroid-myometrium interface were identified: (*i*) sharp, (*ii*) gradual, and (*iii*) absent (Fig. 5c). These categories were quantitatively defined based on thresholds of Δ*E*_*int*_ values (*<* 2 or ≥ 2) and whether fibroids were stiffer than paired adjacent myometrium in a statistically significant manner when all indentation points were considered (Table 3). Representative heat maps of material properties across the fibroid-myometrium interface are shown in Fig. 5f alongside matching brightfield microscopy images. Compiled spatial data for all samples will be made available on Columbia University’s Academic Commons (link provided upon paper acceptance). Of the 34 fibroids assessed in this manner, 11 sharp (32.4%), 19 gradual (55.9%), and 4 absent (11.7%) interfaces were identified. Fibroids with sharp interfaces were stiffer and less permeable than those with gradual interfaces (Figs. 5g, S19). Differences in elastic modulus were also found between fibroids with sharp and absent interfaces (Fig. 5g). No differences in viscoelastic ratio and diffusivity parameters were noted for the different interface categories (Fig. S19). Fibroid size differed slightly between sharp and absent interface categories (*p* = 0.022), but not between gradual and sharp categories (Fig. 5j).

**Table 3.** Metrics defining elastic modulus interface categories.

| Interface Category | $\Delta E_{int}$ | $\Delta E_{all}$ | $p < 0.05$ |
| --- | --- | --- | --- |
| Sharp | $\geq 2$ | $\geq 1$ | Yes |
| Gradual | $< 2$ | $\geq 1$ | Yes |
| Absent | — | — | No |

K-means clustering was then applied to the dataset to further evaluate spatial patterns not previously identified with the aforementioned analysis. Two distinct clusters were discovered for the intrinsic permeability (*k*) parameter, revealing differences across the entire adjacent myometrial–fibroid testing region: fibroids in cluster 0 were less permeable than adjacent myometrial tissue, whereas fibroids in cluster 1 exhibited greater permeability (Fig. 5d,h; Fig. S20). These permeability clusters did not vary with fibroid size (Fig. 5j).

Considered together, the identified elastic modulus and permeability clusters were significantly associated with one another as determined by a chi-squared test (*χ*^2^ = 16.99, df = 2, p = 0.0002). Sharp interfaces exclusively overlapped with permeability cluster 0, occurring at a frequency greater than that expected by chance (Figs. 5e). Approximately 70% of the samples with absent or gradual elastic modulus interfaces overlapped with permeability cluster 1 (Figs. 5e).

### Structural and Compositional Properties

Uterine fibroids were found to be primarily composed of collagen fibers and, to a lesser degree, smooth muscle tissue, based on histological assessment of Masson’s Trichrome stains for a subset of mechanically tested samples (Fig. 6). In contrast, distant myometrium tissue largely consisted of smooth muscle cells, organized into fascicles, between which collagen fibers are interspersed (Fig. 6). Overall, based on semi-quantitative image analysis, there was a statistically significant difference in the area fraction ratio of collagen to smooth muscle between tissue types, with fibroids exhibiting an increased ratio of collagen to smooth muscle content compared to distant myometrial tissue (Fig. 6b). There was also a statistically significant difference in the CV of this parameter between tissue types, suggesting fibroids exhibit more intra-sample heterogeneity in terms of composition relative to the distant myometrium (Fig. S21). Variation in fibroid composition was also noted across distinct patients (Fig. S21). Interestingly, the area fraction ratio of collagen to smooth muscle content did not correlate with any material property changes in this dataset (Fig. S22). Further, on a compositional basis, uterine fibroids exhibited decreased hydration (p = 0.0053) compared to patient-matched myometrial tissue (Fig. 6e). The hydration levels of fibroids and myometrial tissue were found to be 84.1 ± 4.0% and 80.0 ± 4.4%, respectively, and were variable across distinct patients (Figs. 6, S24).

**Figure 6.**
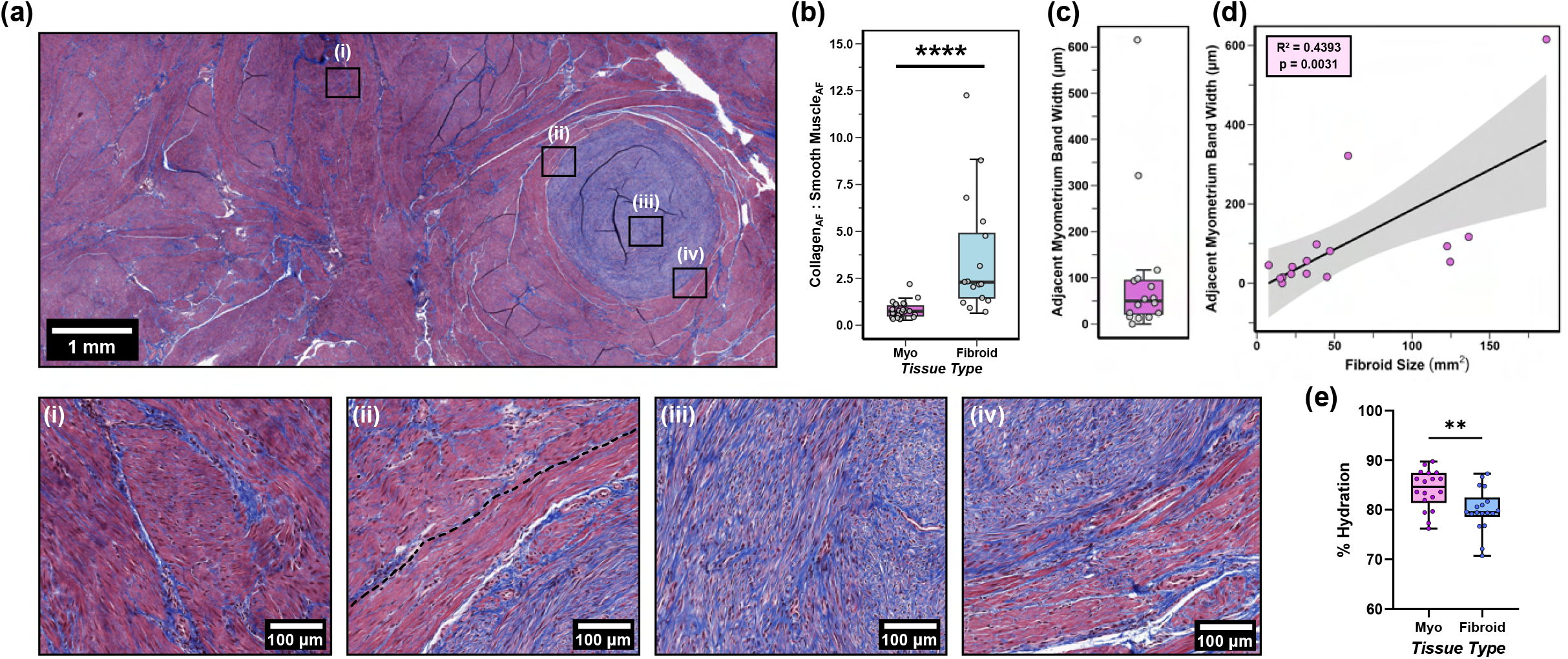
Altered myometrial fiber alignment demonstrated at the seedling fibroid-myometrium interface. **(a)** Representative seedling fibroid (*NP9*.*F2*) and surrounding myometrial tissue stained with Masson’s Trichrome (blue = collagen; red = muscle fibers, cytoplasm; black = nuclei). Selected regions of interest are shown for the (i) distant myometrium, (ii, iv) fibroid-myometrium interface and (iii) fibroid. A band of aligned adjacent myometrial fibers is visualized in (ii) and absent in (iv). The dashed line marks the exterior border of the adjacent myometrium band seen in (ii). **(b)** Quantification of collagen to smooth muscle area fraction (AF) ratios for fibroid and myometrium tissues. Each symbol represents the median value of technical replicates for a distinct sample. Statistical analysis was performed using a linear mixed model, where ^****^ *p <* 0.0001. **(c)** Distribution of adjacent myometrium band widths across samples. Each data point represents the median value of all measurements taken for a distinct sample. **(d)** Adjacent myometrium band width plotted against fibroid cross-sectional area. *R*^2^ and *p* values are noted from linear regression analysis. **(e)** Hydration of myometrium and seedling fibroid tissues. Each dot represents a single measurement taken for a distinct tissue sample. Statistical significance was computed with an unpaired, two-tailed t-test, where ^**^ *p <* 0.01).

Visual assessment of collagen fibers with second harmonic generation (SHG) imaging revealed multiple fiber directionalities for individual fibroids (Fig. 7, S26). The organization of collagen fibers was distinct for each sample evaluated, exhibiting the canonical whorled and mixed phenotypes^34^, with a dense arrangement of collagen fibers (Fig. 7, S26). In contrast, distant myometrial tissue possessed collagen fibers that were less densely arranged and had mixed fiber directionality (Fig. 7). Elastin content appeared both diffuse throughout each tissue type and localized to blood vessels (Fig. 7, S26). Mean signal intensities measured from SHG images demonstrated increased collagen content in fibroids relative to the myometrium, whereas elastin intensity did not differ between tissues (Fig. 7). In addition, the collagen-to-elastin intensity ratio was elevated in fibroids relative to myometrial tissues (Fig. S25).

**Figure 7.**
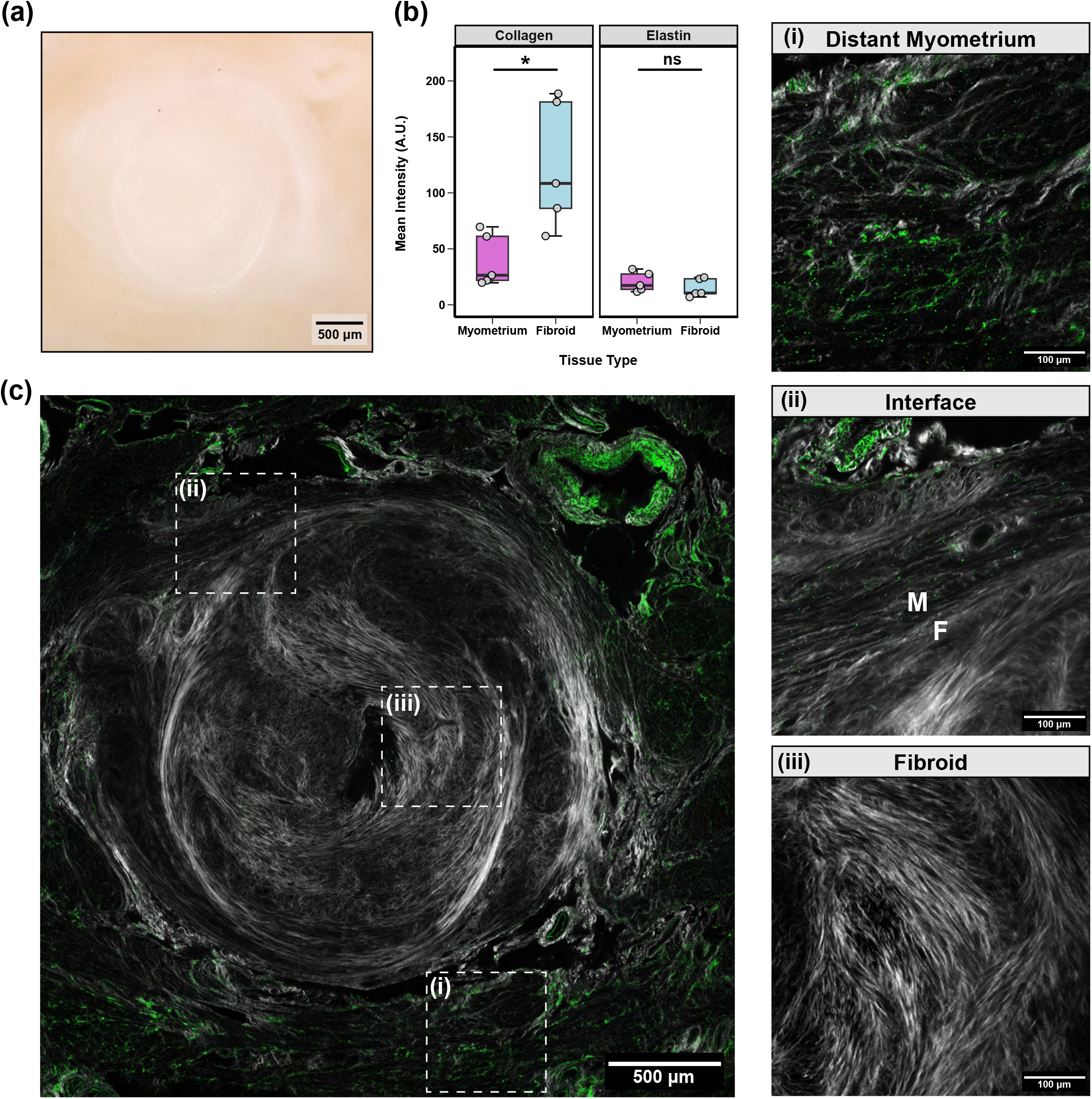
Distinct collagen and elastin fiber organization revealed for seedling uterine fibroids and patient-matched myometrium. **(a)** Brightfield image of a representative seedling fibroid (NP1.F5) embedded within myometrial tissue with **(c)** corresponding second harmonic generation (SHG) imaging. Images are shown for a single imaging plane with collagen (*white*) and elastin (*green*) signals overlaid. Selected regions of interest are shown for **(i)** distant myometrium tissue, **(ii)** the fibroid-myometrium interface (M = myometrium, F = fibroid), and **(iii)** fibroid tissue. **(b)** Quantitative SHG image analysis for myometrium and fibroid tissues in the collagen and elastin channels. Mean intensity with arbitrary units (A.U.) is shown. Each dot represents the mean value of technical replicates for a given sample. Statistical analysis was performed with a Welch’s t-test with a Bonferroni correction with significance marked as follows: ^ns^ *p >* 0.05, ^*^ *p <* 0.05).

### Structural Assessment of the Fibroid-Myometrium Interface

A band of aligned myometrial smooth muscle fibers around the circumference of the fibroid was observed in a large number of histological specimens stained with Masson’s Trichrome (Figs. 6, S28). The width of this band was quantified (median: 56.3 *µ*m), ranging in values from 0 *µ*m (i.e., nonexistent) to 798 *µ*m (Figs. 6, S28). Even for an individual fibroid, a large degree of heterogeneity in the appearance of adjacent myometrial tissue was observed as this band was not always visible along the entire circumference of the fibroid, and in some cases, the interface was wholly unclear (Figs. 6, S28). Differences in this width measurement were also noted for different histology slice depths, suggesting that the plane in which this phenomenon occurs varies (Fig. S28). Interestingly, a positive linear relationship between the width of the adjacent myometrium band and fibroid size was observed; as fibroid size increases, so does the width of the adjacent myometrium band (Fig. 6d). This alignment of myometrial fibers was also visualized with SHG imaging (Fig. 7c,ii).

## Discussion

This study presents a comprehensive spatial atlas of the mechanical, structural, and compositional properties of seedling uterine fibroids at the fibroid-myometrium interface. This novel structure-function dataset, comprising more than 12,000 individual mechanical measurements, represents an extensive characterization of the micro-mechanical properties of human seedling uterine fibroids and myometrium tissues to date. Altogether, seedling uterine fibroids exhibited increased tissue stiffness, decreased permeability, increased diffusivity, increased collagen content, distinct collagen fiber organization, and decreased hydration compared to patient-matched myometrium tissues. Uterine fibroids also exhibited larger degrees of heterogeneity, both mechanically and compositionally, within and across distinct fibroids. These findings suggest that uterine fibroids exhibit unique biophysical fingerprints that differ from those of adjacent and distant myometrial tissues. Through spatial analysis, multiple mechanical interface patterns were identified in this study; elastic modulus values exhibited sharp, gradual, and absent changes between fibroid and myometrium tissues, and two distinct permeability clusters were identified.

Overall, seedling uterine fibroids were found to be stiffer than patient-matched myometrium by 2 − 3 fold in this study (Fig. 2a). This relative mechanical difference is comparable to measurements previously reported in the literature that have evaluated centimeter-sized uterine fibroids and patient-matched myometrium tissue with indentation, compression, and shear^48,51,52,57–59^. However, a subset of samples reported in this dataset and elsewhere in the literature^52^ diverges from this trend. In this study, one fibroid specimen showed no change in stiffness compared to myometrial tissue, and one fibroid sample (NP8.F3) exhibited a decrease in stiffness relative to the adjacent myometrium (Fig. S6). Similarly, Purdy et al. (2020) reported two fibroids with reduced stiffness compared to patient-matched myometrium, which was attributed to tissue degeneration^52^. Although degeneration was not noted for any samples in our dataset, these findings underscore the heterogeneous nature of uterine fibroids, and further investigation into the variable presentations of uterine fibroids, including degenerated, calcified, pedunculated, and parasitic fibroids, is needed.

Compared to values reported in the literature^34,48,51,57,59,60,69^, stiffness measurements of myometrial and fibroid tissues appear to be collectively lower in our dataset. This systemic attenuation of stiffness reported in this study may be a product of tissue processing methods or the mechanical testing approach employed. With regard to tissue processing, multiple freeze-thaw cycles have the capacity to alter the mechanical properties of biological tissues^70,71^, though effects are not always observed^61,72^. Additionally, under indentation, sample thickness is an important consideration to avoid substrate-induced stiffening effects^73^. As such, care was taken to ensure that all tissue specimens were at least 1 mm thick and tested away from the specimen’s edge to avoid introducing any substrate or edge effects. From a mechanical testing approach, the testing method (e.g., indentation, shear, tension), sample configuration, and the strain induced during loading can have a significant effect on the calculated material parameters^74^. The mechanical properties of uterine tissue have been previously shown to be strain and frequency-dependent, exhibiting tension-compression asymmetry^59,61^. This study measured the micro-mechanical properties of uterine tissues within the linear elastic regime (∼ 5% strain) under indentation (R = 50 *µ*m), which subjects samples to a complex loading profile of compression, radial tension, and shear^75^. Modulating mechanical testing parameters, such as indentation depth, probe radius, and prescribed ramp time, can dramatically affect the measured material properties of uterine tissue^75–78^. Microindentation has only been utilized by one other study in the literature for uterine fibroid characterization^52^, but notable differences in testing parameters, including the probe size (R = 2.5 *µm*), tissue thickness (10 *µ*m), and indentation depth (not reported), preclude direct comparison to the dataset of this study. Therefore, future studies are needed to systematically determine the effect of freeze-thaw effects on uterine tissues and evaluate the mechanical properties of these tissues under different mechanical testing modalities.

Time-dependent material behavior is another important consideration in the study of uterine fibroids, as these properties are able to drive abnormal cell behavior independent of stiffness^56^. This study evaluated the combined time-dependent response of poroelasticity and viscoelasticity, with the utilization of the PVE model, to determine measures of intrinsic permeability, viscoelastic ratio, and diffusivity. On the scale of microindentation, uterine fibroids were found to be less permeable and more diffusive than myometrial tissues (Fig. 2a). Still, permeability varied on an individual fibroid basis, and decreased permeability was exhibited by most (70%) but not all specimens (Fig. S6). Notably, k-means clustering successfully identified two spatial patterns in permeability values across the fibroid-myometrium interface, describing the relative increase or decrease in this metric across tissue types (Fig. 5d). Interestingly, no change in viscoelastic ratio was observed across all tissue types evaluated in this study (Fig. 2a). Viscoelastic ratio is a measure that reflects the degree of energy dissipation and describes how elastic or viscous a material is, exhibited no difference across fibroid and myometrium tissues in this dataset (Fig. 2a). Similar measures of energy dissipation (i.e., tan *δ*) reported for fibroid and myometrium tissues under compression also show no difference between tissue types^57,59^. This finding is rather surprising given the underlying structural and compositional differences between these two tissue types. However, this lack of difference in tissue viscoelasticity may be more reflective of the mode of tissue deformation. Under compression, time-dependent behavior is primarily governed by poroelasticity due to fluid pressurization and engagement of the nonfibrillar ground substance (e.g., glycosaminoglycans and proteoglycans)^79–81^. Although the fiber network is unable to sustain compressive loads, it does constrain the lateral expansion of the ground matrix^79^. In contrast, under tension, viscoelasticity dominates when the fiber network is load-bearing^79–82^. Therefore, differences in the viscoelastic behavior of uterine fibroids and myometrium tissues may only become apparent under tensile loading.

It is interesting to highlight the unique relationship between elastic modulus and intrinsic permeability for uterine fibroids, which was not demonstrated by either adjacent or distant myometrium tissue; as elastic modulus increases, permeability decreases (Fig. 2c). This indicates that increased tissue stiffness is correlated with reduced interstitial fluid flow within fibroid tissues due to decreased pore size. Pore size has also been shown to modulate cell ingrowth, vascularization, and nutrient diffusion within biomaterials^83^, and decreased pore sizes in fibroid tissues may be responsible for further driving its pathology (Fig. S5). Additionally, this distinct relationship may offer useful insights into tissue characteristics and serve as a unique biophysical fingerprint for fibroids, reflecting the distinct cellular and extracellular matrix (ECM) composition of this tissue. A previous paper published by our group utilized similar microindentation testing techniques to map the material properties of non-human primate uterine tissue layers and found that the perimetrium exclusively exhibited a negative linear relationship between elastic modulus and intrinsic permeability on a log-log scale^84^. In both datasets, no relationship between elastic modulus and permeability was observed for myometrial tissues, regardless of pregnancy state or distance to fibroid pathology^84^. Since collagen is the predominant ECM component of both perimetrium^45,84^ and uterine fibroid tissues^34,47,49^, we therefore posit that collagen density and cross-linking drive this unique relationship.

Notable spatial mechanical heterogeneity was observed for the fibroids evaluated in this study (Fig. 5). Within the tested region, maximum values of elastic modulus were localized to fibroid tissues, which were equally distributed relative to the fibroid’s center (Figs. 5b, S15). Local tissue stiffness may be an important driver in disease progression and can lend insight into the nucleation and growth patterns of these tissues. Within solid cancerous tumors, local sites of high tissue rigidity have been shown to contribute to breast cancer invasiveness and promote tumor growth^85–87^. Interestingly, the stiffest points identified within the tested fibroid region do not cluster towards the tissue’s center (Fig. 5b). Potentially, this suggests that tissue growth is not solely initiated in the center of the fibroid, as initially hypothesized, but at multiple locations including the peripheral edge. Future studies are needed to investigate the active dynamics of uterine fibroid growth from a mechanical purview.

Understanding the mechanical transition patterns that occur at this interface is relevant for improving our fundamental understanding of fibroid pathophysiology and advancing clinical detection tools for margin identification. In the literature, mechanical interface investigations have largely focused on orthopedic tissues, such as the tendon-bone enthesis and the osteochondral interface^88^. Orthopedic tissue interfaces often exhibit mechanical gradients between dissimilar tissues, exhibiting orders of magnitude differences in elastic moduli to minimize stress concentrations in the transfer of loads^88,89^. These gradients can be gradual or sharp and can modulate tissue growth, nutrient transport, and cell-cell signaling at tissue boundaries^88^. Whether the transition between uterine fibroids and myometrial tissue exists as a gradient or sharp discontinuity has direct implications on the fluid migration patterns between these two tissue types, which can ultimately drive disease progression^90,91^. Further, mechanical gradients can alter drug delivery efficacy and may aid in the prediction of fibroid responsiveness to certain therapies^92,93^. This study specifically sought to investigate the mechanical gradients that occur at the interface of uterine fibroid and myometrium tissues and identified distinct spatial patterns for elastic modulus and permeability parameters (Fig. 5). For elastic modulus, interfaces were categorized as sharp, gradual, or absent depending on the relative changes at the interface and across the entire tissue (Fig. 5c). Two distinct permeability clusters were also identified in this study, varying in the degree of fibroid permeability relative to the adjacent myometrium (Fig. 5d). Overall, this characterization scheme offers a novel approach to quantifying the fibroid-myometrium interface and can serve as a starting point for categorizing fibroid subtypes based on functional metrics. However, it does not capture the full heterogeneity of all fibroid variants, and further refinement or expansion may be necessary in the future. It is important to note that the samples evaluated in this study represent a particular length scale (*<* 1 cm in diameter) and trends could vary when larger fibroids are mapped in a similar manner. When interpreting the results of this study, it is also important to consider the choice to characterize a single interface between fibroid and adjacent myometrium tissues (Fig. 1c). It is entirely possible that a single fibroid may exhibit multiple transition patterns along its circumference and at different tissue planes. A single fibroid (NP2.F9) in this dataset was mechanically mapped along two orthogonal fibroid edges; both interfaces exhibited gradual patterns. While this observation is interesting to note, it is by no means a statistically powered finding. Additional studies are needed to thoroughly investigate this question, ideally expanding this characterization to include all portions of fibroid tissue, rather than just a rectangular strip.

Structural heterogeneity of cell and ECM components was observed for seedling uterine fibroids in this study (Figs. 6, 7). The ratio of collagen to smooth muscle content notably varied across distinct fibroids (Figs. 6b, S21). SHG imaging and histological analysis also revealed unique spatial patterns of collagen fibers within each fibroid, suggesting the existence of multiple morphological phenotypes (Figs. 7, S26). This variability at the seedling stage may underlie the architectural patterns observed in centimeter-sized fibroids, as reported by Jayes et al. (2019)^34^. Additionally, the fusion of multiple fibroids, as thought to be observed for NP1.F8, could be another contributor to the heterogeneous architectural patterns found for larger fibroids^34^. Notably, none of the quantitative structural measures evaluated in this study correlated with the reported mechanical properties (Fig. S22). Similarly, Jayes et al. (2019) reported that fibroid stiffness did not correlate with overall collagen content^34^. This suggests that structural and compositional features not evaluated in this study, such as collagen cross-linking and glycosaminoglycan content, may instead drive the mechanical differences demonstrated for fibroid and myometrial tissues.

Structural and compositional features at the fibroid-myometrium interface suggest that growing fibroids alter the directionality of muscle fibers for the adjacent myometrium (Figs. 6, 7c). As fibroid size increases, so does the band of aligned myometrial tissue adjacent to the fibroid (Fig. 6d). This collective fiber alignment likely arises from growth and remodeling processes driven by active cell responses and passive tissue displacement. Uterine fibroids, which exist as stiff inclusions embedded in a soft tissue matrix, can impart mechanical stresses onto the surrounding myometrium^94–96^. Such forces can trigger the cells in the adjacent myometrium to alter their orientation perpendicular to the direction of strain, similar to what has been demonstrated *in vitro*^47,96^. In this way, active remodeling of the adjacent myometrium at a cellular level could act as a stress-shielding mechanism to prevent tissue damage. Another possibility is the passive displacement of adjacent myometrium tissue due to rapid fibroid tissue growth. As fibroids grow, accumulating greater tissue mass, surrounding smooth muscle tissue can undergo stretch, which in turn contributes to fiber realignment perpendicular to the radial direction of fibroid growth^97^. The speed at which fibroids grow likely dictates the relative contribution of active and passive tissue remodeling and is an important avenue for future investigations.

Several methodological limitations should be considered when interpreting the findings of this study. While uterine fibroids are inherently three-dimensional structures, this study only examined a single two-dimensional cross-sectional tissue plane. Given that samples were microtomed to ensure a flat surface for testing, it is unknown whether and to what extent tissues were evaluated offset from their midplanes and how this potential offset could affect the material and structural properties evaluated in this dataset. The method of fibroid edge detection could also be a potential source of error. Identification of the fibroid edge was done visually when the probe was directly in contact with the tissue. Even though this was performed by a skilled user of the microindenter (D.F.), inherent error is associated with this approach, which could contribute to a skewed identification of the interface and, thereby, the calculation of interface fold changes. While microindentation is a powerful tool to measure the time-dependent material properties of biological tissues spatially, it does not come without its limitations and drawbacks. From a technical sense, the Piuma microindentor, which relies on optical interferometry and sensitive feedback mechanisms, does not always precisely match the prescribed indentation depth of the test and can begin its indentation protocol while the probe is already slightly in contact with the sample (∼ 1 *µm*). Such technical errors can skew the actual strain experienced by the tissue and cause pre-loading of the specimen, contributing to slight errors in the material properties measured. The choice of material model can also have a drastic effect on the fitted material parameters. In this study, the combined effect of poroelasticity and viscoelasticity was considered with the implementation of the PVE model, which employs an analytical, semi-phenomenologic fit of the load relaxation data^74,77,78^. Alternative approaches that employ more simplistic, empirical or complex, microstructurally-driven mechanical models on the same dataset can yield different results^64,67^ and is thus an important consideration for mechanical analysis.

Patient heterogeneity is another important consideration for this study. Many patients in our study cohort presented with co-morbid pathologies (e.g., adenomyosis, endometriosis, prolapse, etc.) and diverse obstetric histories, as noted in Table S1. Therefore, the myometrium tissue, regardless of its distance away from fibroid tissue, may not be entirely free of pathology. There is some evidence to suggest that the existence of fibroids, regardless of their proximity to myometrial tissue, can affect the vascular network and transcriptomic profile of the myometrium^98,99^. No myometrial tissues in this dataset came from patients entirely free of leiomyomata, and it is unknown whether the mere existence of fibroids alters the material properties of distant myometrium tissue. Yet, it is important to note that we did not observe any significant mechanical differences between adjacent and distant myometrial tissues in this dataset (Fig. 2). Therefore, if uterine fibroids affect the mechanical properties of myometrial tissue, that effect appears to be systemic for an individual patient. Still, large, prospective studies with diverse cohorts are needed to tease out the effect of co-existing gynecological pathologies, parity, age, and hormone status.

Despite the aforementioned limitations, the rich spatial data presented in this study yield key insights into the mechanical nature of seedling uterine fibroids. Seedling uterine fibroids represent a crucial stage in fibroid development and offer a critical window for investigating disease progression. This discovery-driven study elucidates key pathophysiological features of uterine fibroids and lays the foundation for future avenues of research. The complex growth and remodeling patterns of uterine fibroids from millimeter to centimeter length scales have yet to be thoroughly studied. Understanding how intrinsic mechanical and structural features of uterine fibroids relate to their growth dynamics and remodeling patterns is an open area of research that could inform the clinical management of this disease. Additionally, insights into mechanical interface patterns could aid drug delivery strategies and advance the development of improved intraoperative margin detection tools for myomectomy procedures.

## Methods

### Tissue Collection & Sample Processing

In accordance with institutional review board (IRB) approval at Columbia University Irving Medical Center (CUIMC), human uterine tissues embedded with fibroids were collected from nonpregnant women (N = 12) undergoing total hysterectomies for noncancerous indications between the ages of 18 and 50, all of whom provided written, informed consent for their participation in this study. Detailed de-identified patient information, including age, race, ethnicity, gravidity, parity, estimated menstrual cycle stage, current hormone treatment, uterine pathology, and obstetric & gynecologic history, is noted in Table S1. Of note, there is a statistically significant difference (Welch’s t-test, p = 0.0073) in age between nulliparous (46.8 ± 1.3 yrs) and parous (41.4 ± 3.4 yrs) individuals (Fig. S1e). Human subjects included in this study have some overlap with previous papers published by our group^45,61,100,101^, as noted in Table S4.

Full-thickness uterine wall tissue specimens, away from sites of visible pathology, were collected immediately following surgery by a member of the research team at the anterior, posterior, and fundus regions. Tissues were immediately flash-frozen on dry ice and stored at -80^°^C until further processing. To identify uterine fibroids for subsequent testing and analysis, thawed uterine tissue specimens were carefully dissected. Fibroids were initially identified by stiff nodules in the dissected tissue via manual palpation by D.F. and were arbitrarily numbered. To confirm the fibroid phenotype, histologically matched samples stained with H&E and Masson’s Trichrome (see “Histology” section) were reviewed by an experienced, board-certified pathologist (X.C.) specializing in gynecologic pathology and cytopathology. Samples that were not classified as fibroids, due to the absence of a clear tissue demarcation with myometrial tissue, were excluded from the final dataset. In total, 38 fibroids were identified for mechanical characterization. Fibroid sub-type, anatomic location, and dimensions are noted in Table S2. To note, the dataset contained the following distribution of fibroids: 6 submucosal (SM), 31 intramural (IM) and 1 subserosal (SS). All fibroid diameters were less than 1 cm. A portion of distant patient-matched myometrium tissue, at least 5 mm away from any visible fibroids, was also dissected for each of the three anatomic regions.

Prior to microindentation testing, bulk tissue specimens containing uterine fibroids and adjacent myometrium were microtomed to achieve a flat surface. This was done to minimize the number of failed indentation points during microindentation testing, particularly at the tissue interface. Samples were embedded in M-1 Embedding Matrix (Epredia, Kalamazoo, MI, USA) to ensure an optimal sectioning temperature and microtomed with a Leica SM2010 R Sliding Microtome at -20^°^C. Distant myometrium samples were not microtomed prior to microindentation testing.

### Microindentation Testing

Spherical microindentation (Piuma, Optics11Life, Amsterdam, NE) was utilized to map the spatial variations in time-dependent material properties of uterine fibroids and patient-matched myometrium. A 50 *µ*m probe radius with a cantilever stiffness of 0.15 – 0.5 N/m was used. In preparation for testing, mm-thick specimens were adhered to a glass dish with superglue (Krazy Glue, Atlanta, GA, USA) to ensure sample stability. Samples were swelled at 4°C overnight in 1X PBS solution supplemented with 2 mM ethylenediaminetetraacetic acid (EDTA) to ensure mechanical measurements were taken at tissue equilibrium. Immediately prior to testing, samples were equilibrated to room temperature for 30 minutes and subsequently tested in Opti-free contact lens solution (Alcon, Fort Worth, TX, USA) to reduce adhesion between the glass probe and sample^102^. For each patient, three samples of distant myometrium corresponding to the three anatomic regions sampled were characterized with at least 50 indentation points (mean = 150), to capture sufficient intra-sample variability. For samples containing uterine fibroids surrounded by myometrial tissue, the fibroid-myometrium interface was tested as a rectangular region to map the local spatial variation of material properties (Fig. 1c). Each rectangular matrix scan contained multiple indentation points, with the distance between each point fixed at 200 *µ*m. The length (*X*) of the testing region was equivalent to the full length of the fibroid (*L*_*F*_) plus a fixed 3 mm of the adjacent myometrium (*L*_*AM*_). The width (*Y*) of the testing region was centered relative to the fibroid’s width (*W*_*F*_) and equal to approximately 0.4\**W*_*F*_ . Exceptions to this testing scheme are noted in Supplementary Methods. The dimensions of each fibroid along the long and short axes were determined with the microindentor. The edge of the sample was identified visually when the probe was directly in contact with the tissue.

Tissues were indented to a fixed depth of 4 *µm* under indentation control, corresponding to a 5% indentation strain and contact area of 380 *µm*^2^. Following a 2 s ramp to the prescribed indentation depth, the probe’s position was held for 15 s to yield a load relaxation curve approaching equilibrium. All tissue specimens were at least 1 mm thick and underwent approximately one to four cycles of freeze-thaw prior to mechanical testing. Following microindentation testing, all samples were imaged on a dissecting microscope (Olympus SZ61, Tokyo, Japan) equipped with an optic ring light attachment (Schott, Wolverhampton, UK) and a Canon EOS Rebel T1i camera using EOS Utility software (Canon, Huntington, NY, USA). Scale bars were then overlaid with ImageJ (NIH, Bethesda, MD, USA). Following microindentation testing, samples were subsequently stored at-80^°^C for use in ancillary imaging studies^101^.

### Microindentation Data Analysis

Using customized MATLAB (Mathworks, Natick, MA, USA) codes, microindentation data was subsequently analyzed with the Hertzian contact model^66^ to determine apparent elastic modulus (*E*_*r*_) in the linear elastic regime and a combined poroelastic-viscoelastic (PVE) model^77^ to determine the following time-dependent material properties: instantaneous elastic modulus (*E*_0_), equilibrium elastic modulus (*E*_∞_), apparent poroelastic modulus (*E*_*PE*_), viscoelastic ratio (*E*_∞_/*E*_0_), intrinsic permeability (*k*), and diffusivity (*D*). When reporting data on fibroid and adjacent myometrial tissues in which the interface region is not the focus of analysis, microindentation data at and around (± 400 *µ*m) the identified interface x-position was removed prior to subsequent analysis, as depicted in Fig. 5a, to account for the curvature of uterine fibroid tissue (Fig. S23). The relative change in material parameters (Δ*E*_*all*_, Δ[*E*_∞_*/E*_0_]_*all*_, Δ*k*_*all*_, Δ*D*_*all*_) was quantified as the ratio of fibroids to adjacent myometrial tissue for each distinct parameter.

#### Hertzian Contact Model

Load versus indentation data from the initial loading portion of the indentation protocol was fit with the Hertzian contact model to determine the apparent elastic modulus (*E*_*r*_) of a material at a given indentation point (Fig. 1e,i)^66,67^. This model assumes contact between a sphere and a half-space for a material that is linear elastic. The following equation was used for this analysis:

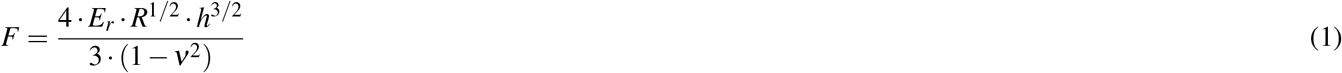

where *F* is the applied force, *R* is the probe radius, *h* is the applied indentation depth, and *ν* is Poisson’s ratio. Here, we assumed tissue incompressibility, setting *ν* equal to 0.5. Data fitting was performed with a customized code in MATLAB using a nonlinear least-squares solver, identical to what has been previously published in the literature^45,77^. Data points were excluded if the corresponding *R*^2^ value was less than 0.5, indicating a poor model fit.

#### Poroviscoelastic (PVE) Model

To determine the time-dependent material properties at a given indentation point, load versus time data from the hold portion of the indentation protocol were fit with a combined poroelastic-viscoelastic (PVE) model in MATLAB based on an established analytical solution with a nonlinear least-squares solver (Fig. 1e,ii)^74,76,77^. The coupled effect of the material’s poroelastic (*P*_*PE*_) and viscoelastic (*P*_*VE*_) force responses is described by:

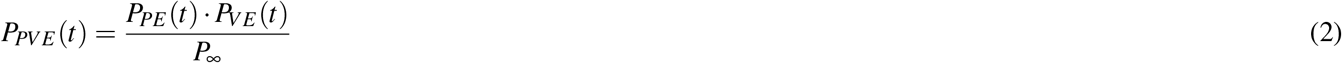

The viscoelastic force response is calculated using a generalized Maxwell model, consisting of a linear spring connected in parallel with two Maxwell units, each containing a linear spring and dashpot connected in series. The viscoelastic component of the model is defined by the following equation:

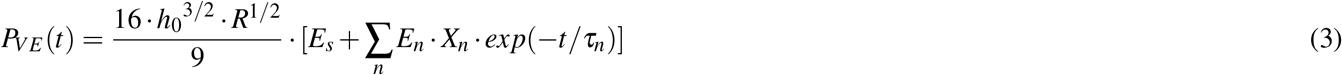

where *E*_*s*_ and *E*_*n*_ are the elastic moduli of the linear spring and the *n*^*th*^ Maxwell element (n = 2), respectively, *h*_0_ is the applied indentation depth, *R* is the probe radius, and *τ*_*n*_ is the characteristic relaxation time of the *n*^*th*^ Maxwell element (n = 2). A ramp correction factor (*X*_*n*_ = (*τ*_*n*_*/t*_*r*_) · [*exp*(−*t*_*r*_*/τ*_*n*_) − 1]) is included to account for the two-second ramp time (*t*_*r*_) since the original Maxwell model assumes a step loading function. Instantaneous elastic modulus (*E*_0_) and equilibrium elastic modulus (*E*_∞_) parameters are determined from Eqn. 3 when *t* = 0 and *t* = ∞, respectively. From these two parameters, viscoelastic ratio (*E*_∞_/*E*_0_), a parameter which describes how elastic (*E*_∞_/*E*_0_ = 1) or viscous (*E*_∞_/*E*_0_ = 0) a material behaves, is calculated.

The poroelastic force response is calculated from the analytical solution published by Hu et al. 2010:

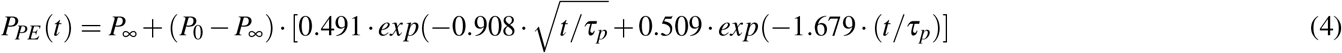

*P*_0_ is the initial force at the beginning of the load relaxation curve and is calculated from the model defined in Hu et al. 2010 as *P*_0_ = (16*/*3)· *G*_*PE*_ · *R*^1*/*2^ · *δ*_0_^3*/*2^, where *G*_*PE*_ is the apparent poroelastic shear modulus. *P*_∞_ is the estimated equilibrium force given by *P*_∞_ = *P*_0_*/*[2(1 − *ν*_*d*_)], where *ν*_*d*_ is the drained Poisson’s ratio. *τ*_*p*_ is the poroelastic time constant defined as *τ*_*p*_ = *a*^2^*/D*, where *a* is the indentation contact radius 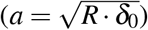 and *D* is diffusivity, a measure that describes the flow of fluid through a porous medium in a one-second period. Intrinsic permeability (*k*), a measure of fluid flow through a porous material, is calculated as follows:

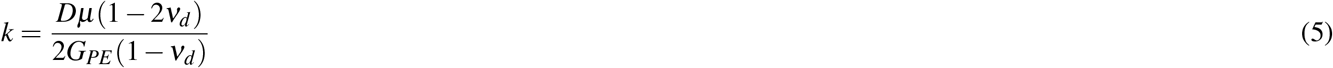

The interstitial fluid viscosity (*µ*) is assumed to be equivalent to the dynamic viscosity of water at 25^°^C (∴ *µ* = 0.89 x 10^−3^ *Pa* · *s*). Material incompressibility is assumed, wherein material volume does not change under applied deformation, and therefore, undrained Poisson’s ratio (*ν*) is set as 0.5. The apparent poroelastic modulus (*E*_*PE*_) is calculated from apparent poroelastic shear modulus as *E*_*PE*_ = 3*G*_*PE*_ . Average pore size (*ξ*) is calculated as the square root of permeability 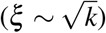 .

Fitted data points were excluded from the final dataset if the load relaxation curve displayed (i) sharp discontinuities, (ii) increasing loads over time, or (iii) ΔP ∼ (*P*_*max*_ – *P*_*min*_) less than 0.01 *µ*N.

### Spatial Analysis of Mechanical Properties

#### Relative Spatial Positioning of Fibroid Extrema

The location (Δ*x*, Δ*y*) of the maximum and minimum elastic modulus (*E*_*r*_) values within the entire tested fibroid region was identified with a custom MATLAB code and was determined relative to the horizontal and vertical midlines of the testing region. Given that the dimensions of the testing region varied with the fibroid dimensions, this relative position was further normalized with respect to the testing region’s length (*X*) and width (*Y*). The normalized position is given by Δ*x*_*norm*_ and Δ*y*_*norm*_.

#### Interface Fold Change Quantification

The relative change in elastic modulus (Δ*E*_*int*_) and intrinsic permeability (Δ*k*_*int*_) was quantified at the fibroid-myometrium interface for all samples containing adjacent myometrial tissue. Relative changes in viscoelastic ratio and diffusivity were not calculated for the interface region due to a lack of statistical significance across tissue types or mathematical dependence on other parameters. Using a custom MATLAB script, the data were first trimmed ± 2 mm around the identified interface x-position. The median value of the adjacent myometrium was then calculated; data points that were within 200 *µ*m of the interface were not included in the calculation. All data were then normalized relative to the median value of the adjacent myometrium region. The data were then trimmed further to only include the center 3 or 4 rows of data for an odd and even number of rows, respectively. This was done to minimize the effect of fibroid tissue curvature in this analysis (Fig. S23). The relative fold change in material parameters was then calculated from the normalized, trimmed data ± 200 *µ*m around the interface x-position.

#### Identification of Interface Patterns

The interfaces of uterine fibroids were grouped into three categories: (*i*) sharp, (*ii*) gradual, and (*iii*) absent. Classification was determined based on the interface fold change of elastic modulus values (Δ*E*_*int*_) and whether fibroids were stiffer than matched adjacent myometrium, as outlined in Table 3. Statistical significance (*p <* 0.05) was determined between the mechanical replicates of matched fibroid and myometrium samples using a Welch’s t-test with a Bonferroni correction.

#### K-Means Clustering Analysis

Using a custom Python script, microindentation data were processed so that all samples were centered on the interface (X0) with the myometrium region on the left (X-) and the fibroid region on the right (X+). Missing data were imputed using the sample median. To ensure confidence in the boundary between uterine fibroid and myometrium tissue measurements and minimize the effect of curvature, the middle value along the Y-axis was used for this analysis (Fig. S27). The final processed data used as input for clustering is included in supplemental materials (Fig. S29). K-means clustering was performed using sklearn’s Kmeans class. To determine the optimal number of clusters, we considered both the elbow and silhouette score (Fig. S30). For intrinsic permeability (*k*), two clusters were chosen (Fig. 5d).

### Histology

Samples were prepared for histology either at the time of initial tissue dissection or following microindentation testing. Samples were fixed in 10% formalin solution for at least 24 hrs, subsequently transferred to 70% ethanol solution, paraffin-embedded, and sectioned to a thickness of 5 *µ*m. To observe gross tissue morphology and the distribution of collagen and smooth muscle, all samples were stained for Hemotoxylin & Eosin (H&E) and Masson’s Trichrome by the Molecular Pathology Core Facilities at CUIMC using standard protocols^103^. Samples were imaged under brightfield microscopy at 20 x magnification with a Leica Aperio AT2 whole slide scanner and visualized with the Aperio ImageScope software (v12.3.1.6002, Leica Microsystems, Wetzler, Germany).

#### Image Quantification

The relative proportion of collagen and smooth muscle content was quantified from Masson’s Trichrome stained samples of fibroid and myometrial tissue. This quantification was performed on a subset of samples prepared at the time of tissue dissection that underwent one to three freeze-thaw cycles. One to five representative images per tissue group were taken at 20x magnification with a Leica DMi1 Inverted Microscope using the Leica Application Suite X (LAS-X). For this quantification, regions containing blood vessels in more than fifty percent of the image area were avoided. The area fractions of blue and red color, corresponding to collagen and smooth muscle content, respectively, were quantified in ImageJ (NIH, Bethesda, MD, USA) with RGB color deconvolution and a thresholding function.

The width of the aligned myometrial fiber band immediately surrounding intramural fibroids was quantified in a subset of samples stained with Masson’s Trichrome; measurements were taken at three to five distinct locations around the fibroid. Measurements were done in either the Aperio ImageScope software for whole scanned slides or ImgaeJ for individual images taken at 10x magnification with the aforementioned inverted microscope. Samples that exhibited the following issues were not included in this analysis: (i) poor Masson’s Trichrome staining, (ii) notable histological artifacts (e.g., folding, gapping, or tearing), or (iii) diffuse appearance of fibers emblematic of excessive tissue swelling.

### Tissue Hydration

Lyophilization was used to determine tissue hydration of uterine fibroid and patient-matched myometrium tissues using a FreeZone 4.5 Liter Benchtop Freeze Dry System (Labconco, Kansas City, MO, USA). Tissues (*mm*^3^) from six patients (NP1, NP2, NP3, NP4, NP5, NP7) were analyzed with this assay. For each individual, three distinct samples of distant myometrial tissue from random anatomic regions were analyzed. Fibroid samples were separated from adjacent myometrial tissue, and the number assessed per patient varied (n = 1–7). Samples included both a subset of fibroids tested previously with microindentation and a group of loose, mechanically untested fibroids. For all samples, wet and dry tissue weights were measured in a pre-weighed 1.5 ml Eppendorf tube before and after lyophilization using an analytical balance (MS105, Mettler Toledo, Greifensee, Switzerland) with 0.01 mg readability. Tissue hydration was calculated with the following equation:

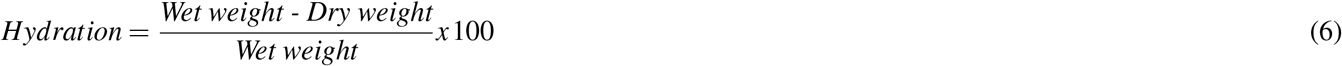

### Second Harmonic Generation (SHG) Imaging

The organization of collagen and elastin fibers in uterine fibroid and myometrium samples was evaluated with SHG imaging. Imaging was performed on a resonant scanning multiphoton confocal microscope (Nikon ECLIPSE Ti2-E inverted microscope equipped with an A1RMP resonant spectral scanning confocal unit and a motorized XY Stage) in the Confocal and Specialized Microscopy Shared Resource (CSMSR) of the Herbert Irving Comprehensive Cancer Center (HICCC) at Columbia University. A Nikon water immersion 25x / 1.1 W Apo LWD objective lens was used for imaging. Artificial tear solution was used in place of water as the immersion media. A subset of mechanically tested samples was imaged with SHG (see Table S3). Prior to imaging, all samples were microtomed flat (see “Sample Processing” section) and swelled overnight in 1X PBS solution. Samples were mounted face-down onto a rectangular (25 x 75 mm) coverglass slide with #1 thickness (Chemglass Life Sciences LLC, Vineland, NJ, USA) for imaging. The Nikon NIS-Elements software was used for image acquisition. Pixel resolution was set as 1024 x 1024; each pixel corresponded to a length of 0.50 *µ*m. X and Y parameters were adjusted based on the unique dimensions of each sample and captured both the fibroid(s) of interest and surrounding myometrial tissue. Each sample was imaged at 3–5 x-y planes, with the distance between each z-stack being 10–15 *µ*m (Table S3). Two excitation wavelengths, 860 nm and 1040 nm, were used to visualize collagen and elastin fibers, respectively^104,105^; the corresponding emission wavelengths were 525 nm and 575 nm, respectively. Laser power was set at 5%. Once images were acquired, multiple images were automatically stitched in the Nikon NIS-Elements software to create overview image of each sample.

#### SHG Image Analysis

All SHG image analyses were performed in ImageJ software (NIH, Bethesda, MD, USA). Regions of interest (ROIs) for fibroids were manually cropped and represented the whole, discernible cross-sectional tissue region. Multiple ROIs of myometrial tissue, away from the identified fibroid margins, were analyzed per image. For both tissue types, measurements were taken in the same ROI location across multiple z-stacks (n = 3–5). For each distinct tissue sample, replicate data across z-stacks and different ROIs were averaged prior to statistical analysis. Mean signal intensity was analyzed for images of fibroids and myometrial tissue taken in the 860 nm and 1040 nm excitation wavelength channels, corresponding to collagen and elastin content, respectively. For each identified region of interest (ROI), mean signal intensity was calculated as total signal intensity divided by ROI area.

### Statistical Analysis

Statistical analysis was performed using RStudio (v2023.09.0), GraphPad Prism (v.10.2.2), and Python (v3.13.0). The normality of all data was first assessed using Q-Q plots. In instances of non-normal data distributions, data were normalized with a logarithmic transformation. A linear mixed-effects model was employed to analyze all datasets in this study unless otherwise noted, with Patient ID or Sample ID set as the random variable, where appropriate. Pairwise contrasts between tissue layers were obtained using estimated marginal means (emmeans) with containment-based degrees of freedom. Hydration data was analyzed with an unpaired, two-tailed t-test. For SHG intensity analysis, unpaired t-tests with Welch’s corrections were used. For correlation analysis between two continuous variables, Pearson correlation coefficients were calculated; *R*^2^ and *p* values are reported for each fit. Principal component analysis (PCA) was performed using scikitlearn on the median material property parameters (*E*_*r*_, *E*_∞_*/E*_0_, *k, D*) for each sample’s tissue type: fibroid, adjacent myometrium, and distant myometrium. Differences between tissue groups were determined to be statistically significant by permanova. To assess the predictive value of each of the reported material properties, a receiver operating characteristic (ROC) curve, which plots sensitivity as a function of [1-specificity], was generated for comparisons between tissue types, and the corresponding area under the curve (AUC) values were computed with the Wilson/Brown method. Associations between categorical variables for interface sharpness (absent, gradual, sharp) and permeability cluster (0, 1) were evaluated using a Pearson’s chi-square test of independence. Expected cell counts were calculated under the null hypothesis of independence, and standardized Pearson residuals were used to identify category combinations occurring more or less frequently than expected.

For all analyses, significance was set at a 95% confidence level. P-value symbols are defined as follows: ^*^ *p <* 0.05, ^**^ *p <* 0.01, ^***^ *p <* 0.001), ^****^ *p <* 0.0001. No significance (*p >* 0.05), if marked, is indicated with *ns*.

## Acknowledgments

Research reported in this publication was supported financially by the National Science Foundation (NSF) Graduate Research Fellowship (DGE-2036197) to D.F., The Iris Fund (K.M. and T.K.), and National Institutes of Health (NIH) under award numbers 1R01HD091153 **(K.M.)**, 1F31HD114449-01 (D.F.), T32DK007647 (A.S.), and T15LM007079 (A.S.). This study also used the Confocal and Specialized Microscopy Shared Resource of the Herbert Irving Comprehensive Cancer Center at Columbia University, funded in part through the NIH/NCI Cancer Center Support Grant P30CA013696. We would like to thank Dr. Jason Fan for assistance with SHG imaging, Elizabeth Vitaro for confirmation of de-identified patient information, Dr. Camilo Duarte-Cordon for thoughtful scientific discussions, and Dr. Joy Vink and Dr. Shuyang Fang for tissue collection of human uterine specimens. The content is solely the responsibility of the authors and does not necessarily represent the official views of the NIH or NSF. Figures for this paper were created with BioRender.com, Adobe Illustrator, and GraphPad Prism under registered academic licenses.

## Author Contributions Statement

D.F., K.M., and M.O. conceived the study. D.F., A.A., and K.M. secured the necessary human tissues for investigation. D.F. performed all mechanical experiments, while D.F., J.J., A.T., and A.S. conducted compositional and structural assays. X.C. evaluated and validated all histological specimens. D.F. and A.S. analyzed the data, and D.F., K.M., A.S., T.K., and M.O. interpreted the results. D.F. and A.S. prepared all figures and visualizations. K.M., M.O., A.A., C.H., and T.K. supervised the research. D.F. and K.M. obtained funding. D.F. wrote the original draft, and all authors reviewed and approved the final manuscript.

## Data and Materials Availability Statement

All data reported in this manuscript have been made available in supplementary materials or on Academic Commons (a link will be provided upon paper acceptance). Codes used for data analysis are available upon request to the corresponding author (KMM).

