## Supplemental Materials for "Spatially Mapping the Mechanical and Structural Properties of the Seedling Uterine Fibroid-Myometrium Interface"

### SUPPLEMENTAL INFORMATION

#### Microindentation Testing

There are miscellaneous exceptions to the microindentation testing scheme described in the main methods section of the manuscript. These exceptions can be visually observed in supplemental materials made available on Columbia University's Academic Commons (link provided upon paper acceptance). There was one case of a loose fibroid sample (NP1.F1) devoid of adjacent myometrial tissue that was characterized by a 10 x 10 grid of points. There are two cases (NP3.F1.2 & NP6.F1) where the tested adjacent myometrium length was less than 3 mm because of the dissected myometrium tissue dimensions. There is one case containing two fibroids (NP1.F3 & NP1.F4) in a single tissue specimen that was tested as a single rectangular grid of points. NP1.F3 & NP1.F4 were separated by 600$\mu$m of myometrial tissue, which was trimmed entirely from all analyses. Adjacent myometrium for this tissue specimen was tested to the left of sample NP1.F3. There was another case containing two fibroids (NP10.F1.1 & NP10.F1.2) that was tested as a single rectangular grid of points, however these fibroids were separated by an undetermined amount of myometrial tissue. In this case, microindentation data was truncated significantly beyond the standard $\pm$400 $\mu$m around the interface to ensure the material properties reported are truly representative of fibroid tissue; adjacent myometrial properties were not reported in this case. There is one case (NP10.F6.2) where the adjacent myometrium region was expanded to 5mm; reported adjacent myometrium values were truncated to exclude a focal region of whitish tissue. Additionally, there was a single fibroid (NP2.F9) that was tested in two different orientations, perpendicular to one another, to evaluate whether the interface properties varied depending on the sample edge tested.

### SUPPLEMENTAL FIGURES & TABLES

**Table S1.** Detailed Patient Information

| Patient ID | No. of Tested Fib. | Age (yrs) | Race | Eth | Gravidity/ Parity | Estimated Menstrual Cycle Stage | Hormone Therapy | Uterine Pathology | Obstetric & Gynecologic History |
|---|---|---|---|---|---|---|---|---|---|
| NP1 | 7 | 47 | Unk | Unk | 2 / 2 | Weakly Proliferative | None listed | Adenomyosis, Leiomyoma, Chronic endometritis | VDx2 |
| NP2 | 3 | 44 | AA | Not Hisp | 2 / 2 | Late Secretory | None listed | Adenomyosis, Leiomyoma | Tubal ligation, VDx1, CSx1 |
| NP3 | 3 | 49 | Unk | Hisp | 0 / 0 | Proliferative | None listed | Leiomyoma, Chronic endometritis | None |
| NP4 | 8 | 46 | White | Not Hisp | 0 / 0 | Proliferative | None listed | Leiomyoma | Left salpingo-oophorectomy |
| NP5 | 1 | 45 | Unk | Hisp | 1 / 0 | Secretory | None listed | Leiomyoma, Endometrial polyps | VTOPSx 1 |
| NP6 | 1 | 41 | White | Not Hisp | 3 / 2 | None – Post-menopausal | Yes – Progesterone | Atrophic endometrium, Adenomyosis, Leiomyoma | SABx1, VDx2 |
| NP7 | 1 | 47 | White | Not Hisp | 0 / 0 | Inactive | None listed | Adenomyosis, Leiomyoma (degenerating) | Unknown |
| NP8 | 1 | 42 | Unk | Hisp | 3 / 2 | Proliferative | None listed | Adenomyosis, Leiomyoma | VDx2 |
| NP9 | 3 | 47 | White | Unk | 0 / 0 | Unknown | Yes – Norethindrone acetate (progstin) | Leiomyoma | LEEPx1, Left salpingo-oophorectomy |
| NP10 | 8 | 38 | Other | Hisp | 1 / 1 | Unknown | Yes – Norethindrone (progestin) | Endometriosis, Adenomyosis, Leiomyoma | CSx1, Tubal ligation |
| NP11 | 1 | 42 | Other | Hisp | 2 / 2 | Inactive | None listed | Leiomyoma, Negative endometrial tissue | CSx2 |
| NP12 | 1 | 36 | Other | Hisp | 3 / 3 | Proliferative | None listed | Prolapse (uterus & vagina), Adenomyosis, Leiomyoma | LEEPx1, VDx1 |

**[KEY]** No. ≡ Number; Fib ≡ Fibroid; Eth ≡ Ethnicity; Unk ≡ Unknown; AA ≡ African American; Hisp ≡ Hispanic; VD ≡ Vaginal delivery; CS ≡ Cesarean section; VTOPS ≡ Vaginal termination of pregnancy; SAB ≡ Spontaneous abortion; LEEP ≡ Loop electrosurgical excision procedure

**Table S2.** List of fibroids tested with microindentation, with corresponding sample characteristics (i.e., anatomic region, fibroid subtype, and size) noted. Fibroid size reports the long and short axis measurements of each sample.

| Patient ID | Sample ID | Anatomic Region | Fibroid Subtype | Fibroid Size (mm x mm) |
|---|---|---|---|---|
| NP1 | F1 | Fundus | Intramural | 6.3 x 5 |
| | F2 | Fundus | Submucosal | 4.45 x 3 |
| | F3 | Fundus | Intramural | 2.7 x 2.6 |
| | F4 | Fundus | Intramural | 2.6 x 2.6 |
| | F5 | Anterior | Intramural | 2.6 x 2.25 |
| | F8 | Posterior | Intramural | 4.2 x 3.4 |
| | F9 | Posterior | Intramural | 4.2 x 3 |
| NP2 | F1 | Anterior | Intramural | 9 x 6.6 |
| | F8 | Anterior | Intramural | 3 x 2.7 |
| | F9 | Anterior | Intramural | 5.25 x 2.8 |
| NP3 | F1.1 | Anterior | Intramural | 3.1 x 2.4 |
| | F1.2 | Anterior | Intramural | 2.8 x 2.6 |
| | F2 | Anterior | Intramural | 4.6 x 3.6 |
| NP4 | F1 | Anterior | Intramural | 5 x 3 |
| | F3 | Unknown | Intramural | 2.4 x 2 |
| | F5 | Unknown | Intramural | 6.2 x 6 |
| | F7 | Posterior | Intramural | 3.2 x 2.4 |
| | F8 | Posterior | Intramural | 3.8 x 2.7 |
| | F9 | Posterior | Intramural | 2.5 x 1 |
| | F10 | Posterior | Intramural | 8.2 x 5.3 |
| | F12 | Posterior | Intramural | 4.5 x 1.8 |
| NP5 | F2 | Anterior | Intramural | 5.2 x 4.55 |
| NP6 | F1 | Posterior | Intramural | 3.6 x 2.6 |
| NP7 | F2 | Posterior | Intramural | 3.5 x 3.5 |
| NP8 | F3 | Anterior | Intramural | 3.4 x 3 |
| NP9 | F1 | Anterior | Intramural | 3.2 x 3.2 |
| | F2 | Anterior | Intramural | 2.5 x 2.1 |
| | F3 | Anterior | Intramural | 2.6 x 2 |
| NP10 | F1.1 | Fundus | Submucosal | 6 x 5.5 |
| | F1.2 | Fundus | Submucosal | 4.3 x 4.3 |
| | F3 | Anterior | Intramural | 6.4 x 6.2 |
| | F4.1 | Anterior | Intramural | 6.8 x 6 |
| | F4.2 | Anterior | Intramural | 3.2 x 3 |
| | F5 | Anterior | Subserosal | 2.9 x 2.4 |
| | F6.1 | Anterior | Submucosal | 5 x 3 |
| | F6.2 | Anterior | Submucosal | 5.35 x 3.16 |
| NP11 | F1 | Anterior | Intramural | 2.9 x 2.4 |
| NP12 | F1 | Posterior | Intramural | 1.85 x 1.3 |

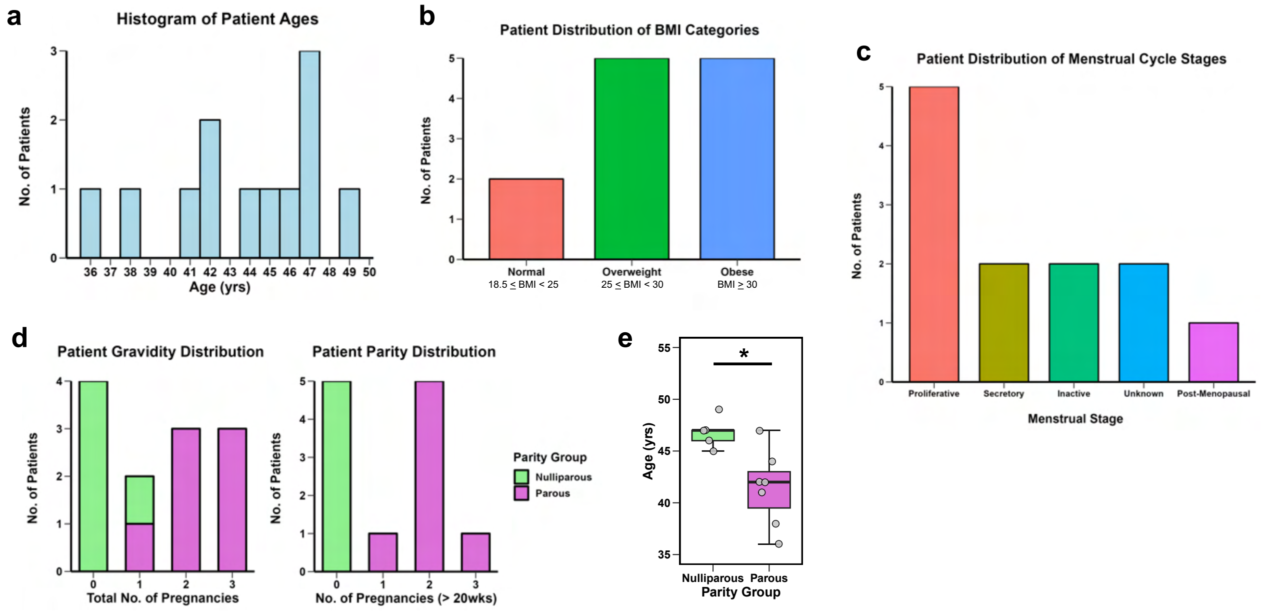


**Figure S1. Summary of patient characteristics.** **(a)** Histogram of patient ages. **(b)** Distribution of body mass index (BMI) categories. **(c)** Distribution of menstrual cycle stages by patient determined from histological evaluation. **(d)** Gravidity and parity distributions by patient. Nulliparous individuals (parity = 0) and parous (parity $\geq$ 1) are colored in green and pink, respectively. **(e)** Differences in age between nulliparous and parous individuals. Statistical significance was computed with a Welch's t-test; $^{*}p < 0.05$.

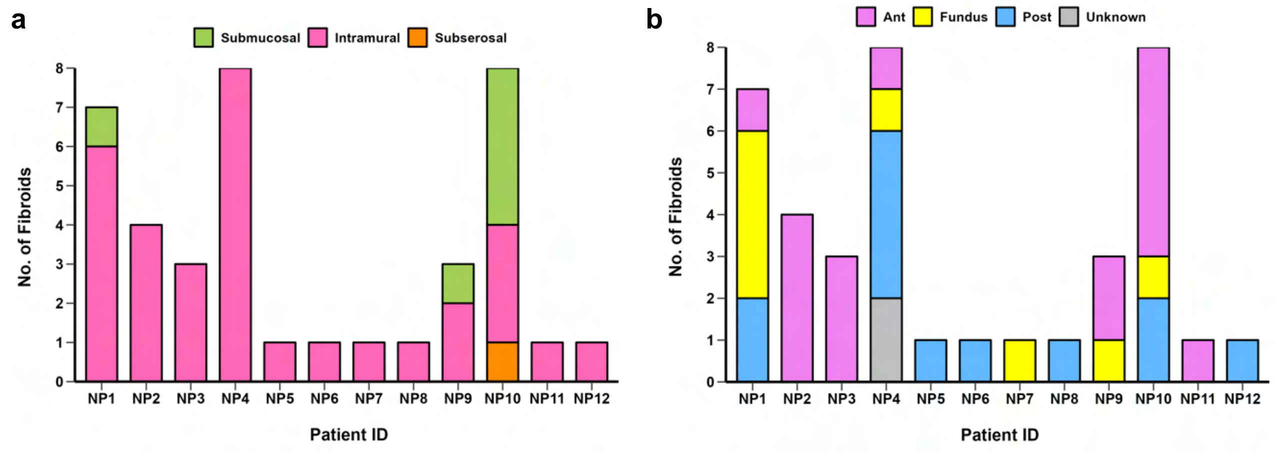


**Figure S2. Distribution of fibroid characteristics by patient.** Data is separated by **(a)** subtype (i.e., submucosal, intramural, subserosal) and **(b)** anatomic region (i.e., anterior, fundus, posterior, unknown).

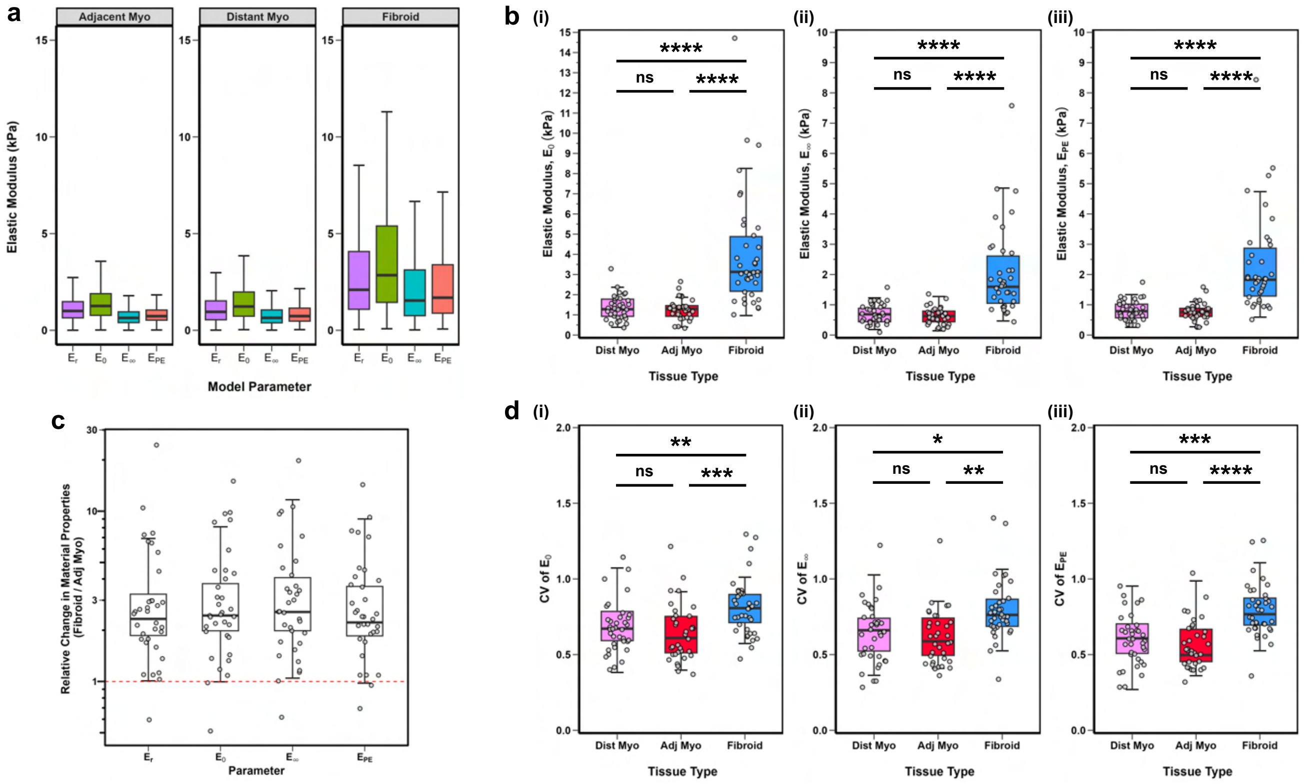


**Figure S3.** **Comparison of all elastic modulus parameters** ($E_r$, $E_0$, $E_\infty$, $E_{PE}$)**.** Absolute values of elastic modulus parameters compared, separated by **(a)** tissue type and **(b)** material parameter. **(c)** Relative fold change in elastic modulus for fibroid samples relative to matched adjacent myometrium tissues. **(d)** Coefficient of variance (CV) for all elastic modulus parameters. Each symbol represents the mean value of all indentation points measured for an individual sample. For all comparison, statistical analysis was performed using a linear mixed model, with sample ID included as a random effect; significance is denoted as follows: $^{ns}$ $p > 0.05$, $^{*}$ $p < 0.05$, $^{**}$ $p < 0.01$, $^{***}$ $p < 0.001$, $^{****}$ $p < 0.0001$.

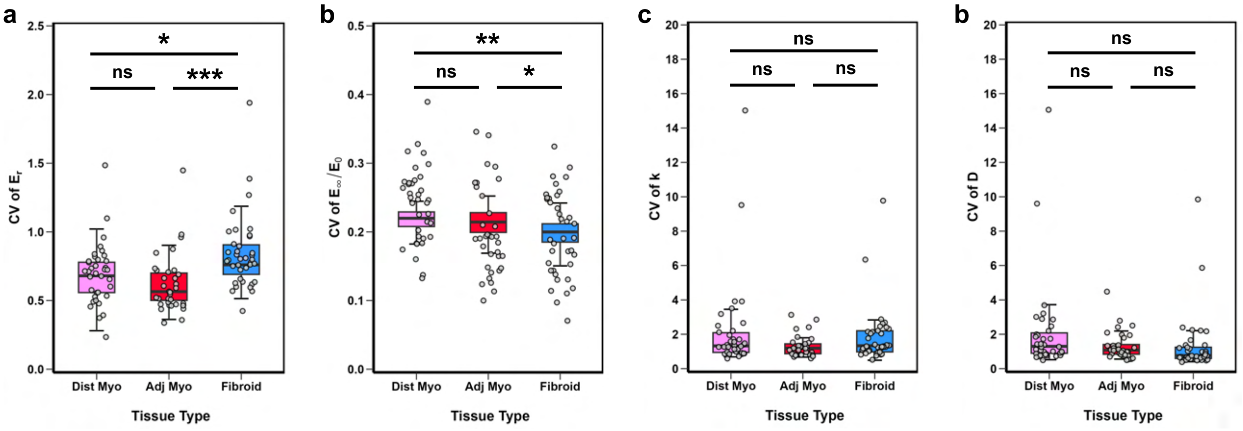


**Figure S4.** **Coefficient of variances (CV) by tissue type.** Data shown for **(a)** apparent elastic modulus [$E_r$], **(b)** viscoelastic ratio [$E_\infty/E_0$], **(c)** intrinsic permeability [k], and **(d)** diffusivity [D]. Each symbol represents the mean value of all indentation points measured for an individual sample. Statistical analysis was performed using a linear mixed model, with patient ID included as a random effect; significance is denoted as follows: $^{ns}$ $p > 0.05$, $^{*}$ $p < 0.05$, $^{**}$ $p < 0.01$, $^{***}$ $p < 0.001$.

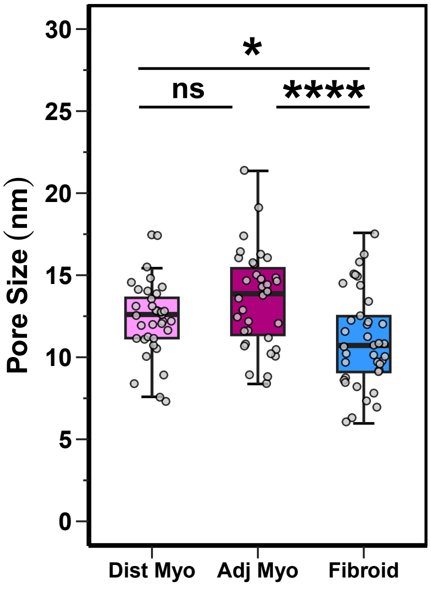


**Figure S5.** **Pore size by tissue type.** Each symbol represents the mean value of all indentation points measured for an individual sample. Statistical analysis was performed using a linear mixed model, with sample ID included as a random effect; significance is denoted as follows: $^{ns}$ $p > 0.05$, $^{*}$ $p < 0.05$, $^{****}$ $p < 0.0001$.

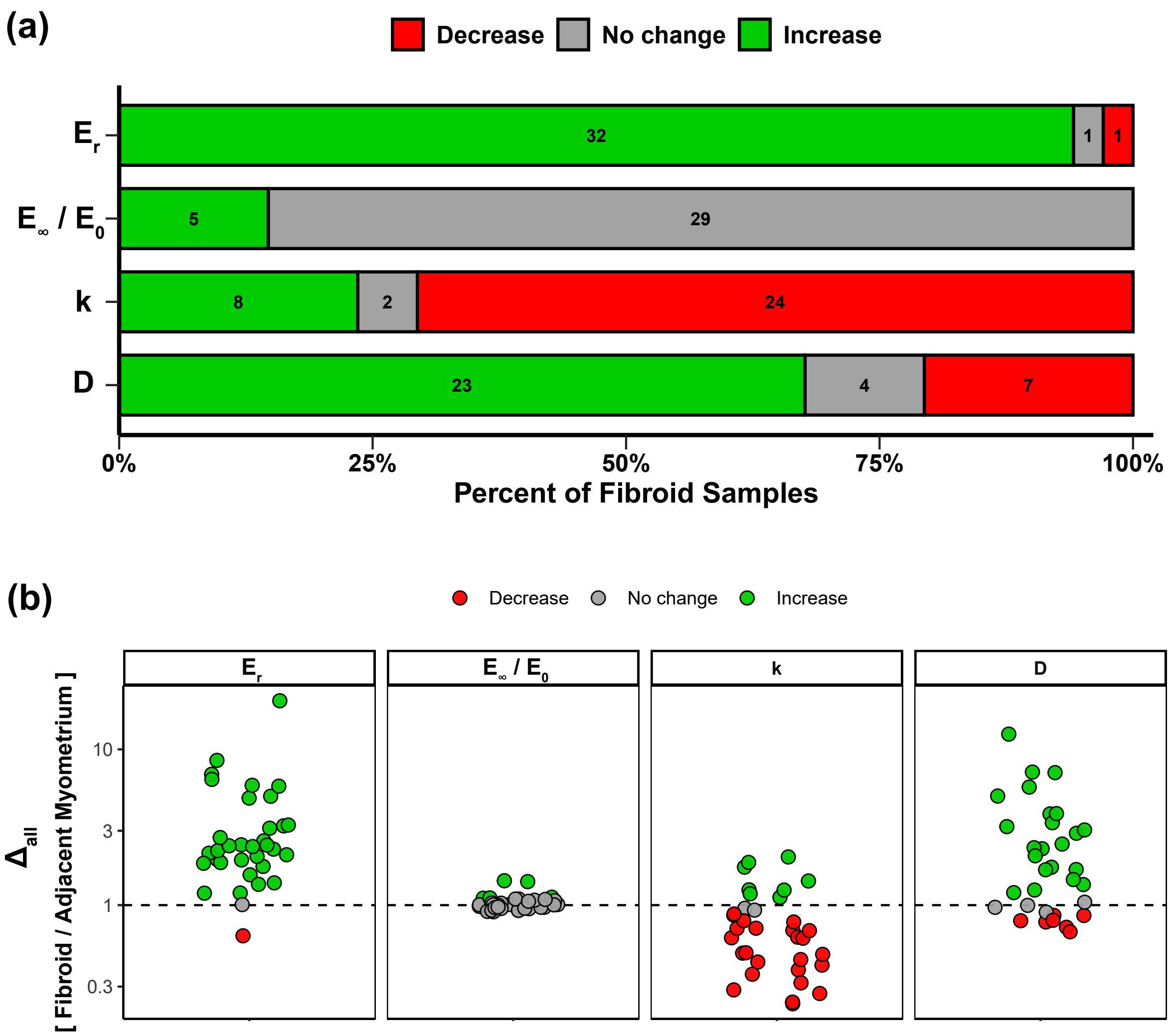


**Figure S6.** **Summary of mechanical heterogeneity across uterine fibroids.** Data show the proportion of fibroid samples that increased ($\Delta_{all} > 1.1$), did not change ($\Delta_{all} = 1 \pm 0.1$), or decreased ($\Delta_{all} > 0.9$) in material properties relative to patient-matched adjacent myometrium. Parameters include the elastic modulus ($E_r$), viscoelastic ratio ($E_\infty / E_0$), intrinsic permeability ($k$), and diffusivity ($D$). .

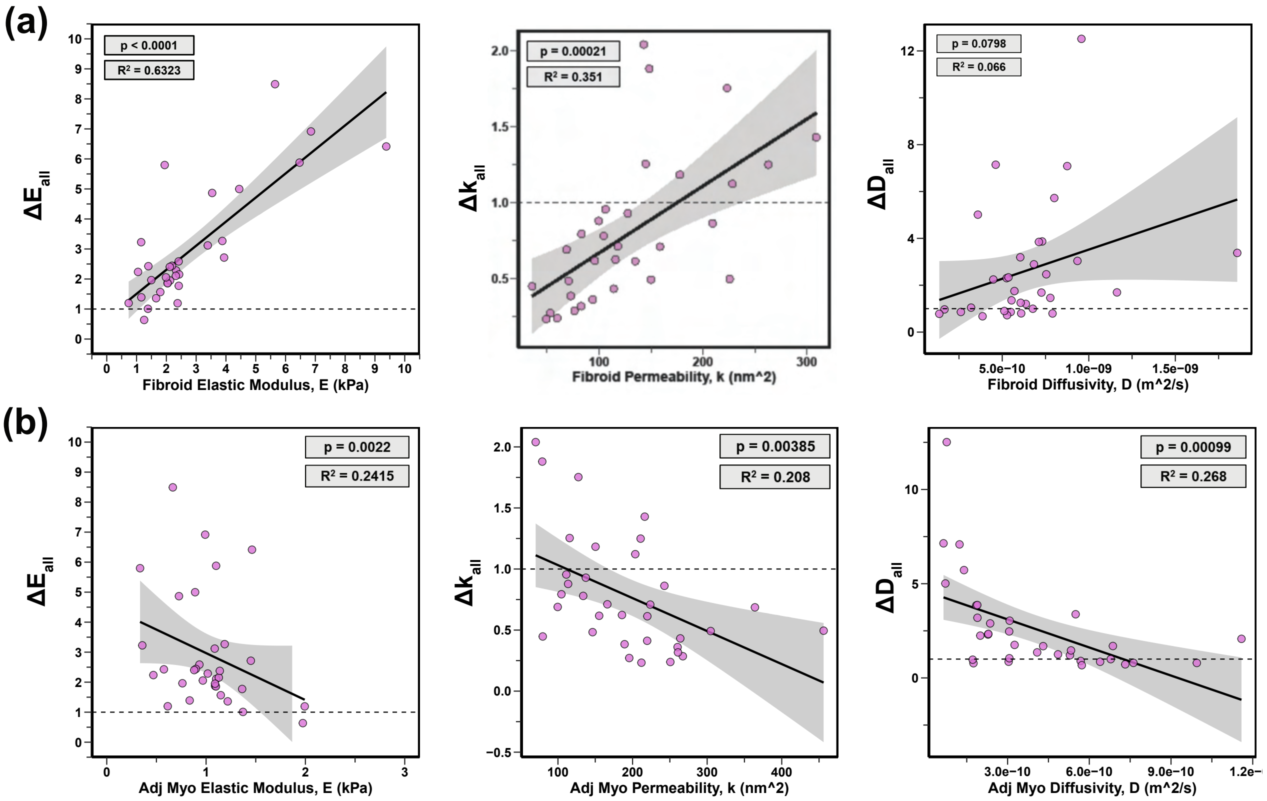


**Figure S7.** Relative change in material properties ($\Delta E_{all}$, $\Delta k_{all}$, $\Delta D_{all}$) plotted versus the absolute value of the corresponding material parameter for (a) fibroid and (b) adjacent myometrium tissues. $p$ and adjusted $R^2$ values are reported for each linear regression analysis. Each symbol represents data for an individual sample. Dashed horizontal line marked at 1.0 indicates no relative change in material properties.

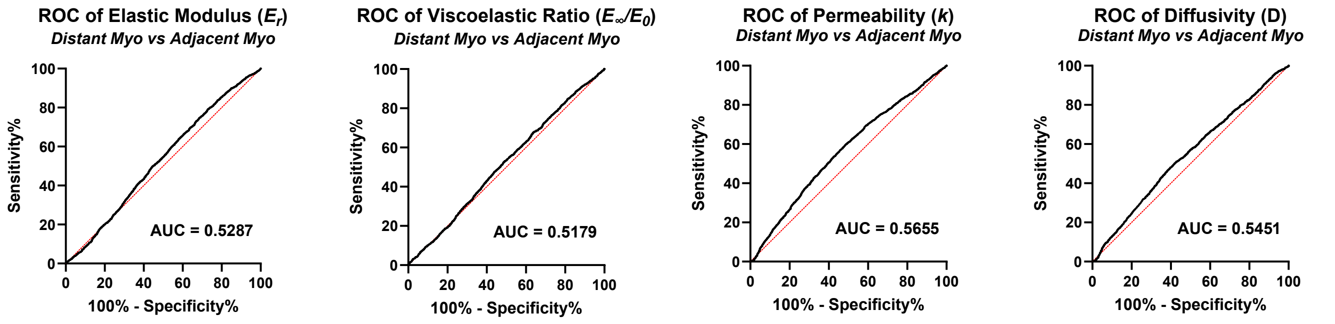


**Figure S8.** ROC curves (*black*) plotting [sensitivity (%)] versus [100% – specificity (%)] between adjacent myometrium and distant myometrium tissues for all four material parameters: elastic modulus ($E_r$), viscoelastic ratio ($E_{inf}/E_0$), diffusivity ($D$), and permeability ($k$). Area under the curve (AUC) values are noted for each comparison. Red lines are included for reference to indicate random chance.

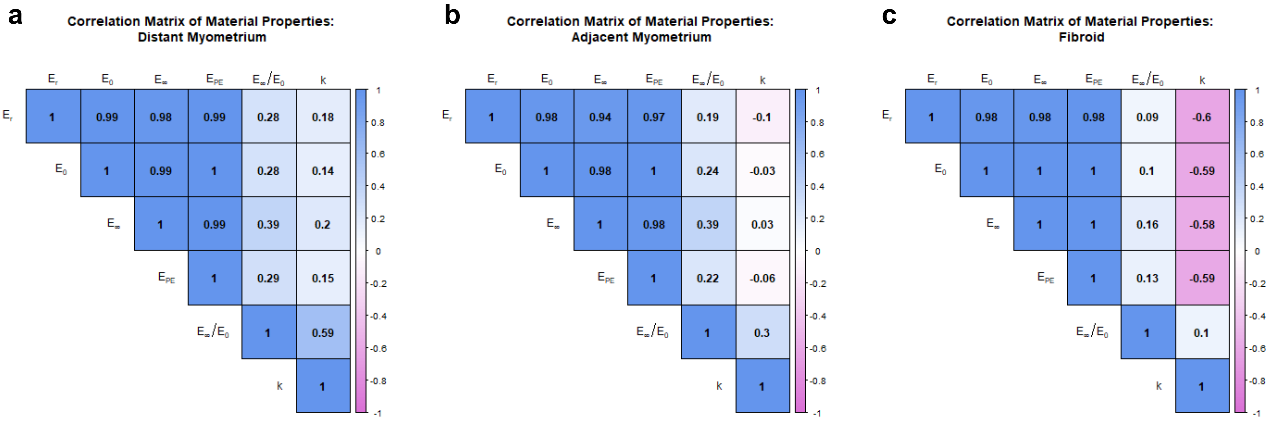


**Figure S9. Correlation matrices of material properties for each uterine tissue type.** Each matrix shows pairwise Pearson correlation coefficients ($r$) among measured material properties for (a) distant myometrium, (b) adjacent myometrium, and (c) fibroid tissues. Material properties include the equilibrium elastic modulus ($E_\infty$), instantaneous elastic modulus ($E_0$), apparent elastic modulus ($E_r$), viscoelastic ratio ($E_\infty/E_0$), and intrinsic permeability ($k$). Positive correlations are shown in blue and negative correlations in magenta, with color intensity proportional to correlation strength.

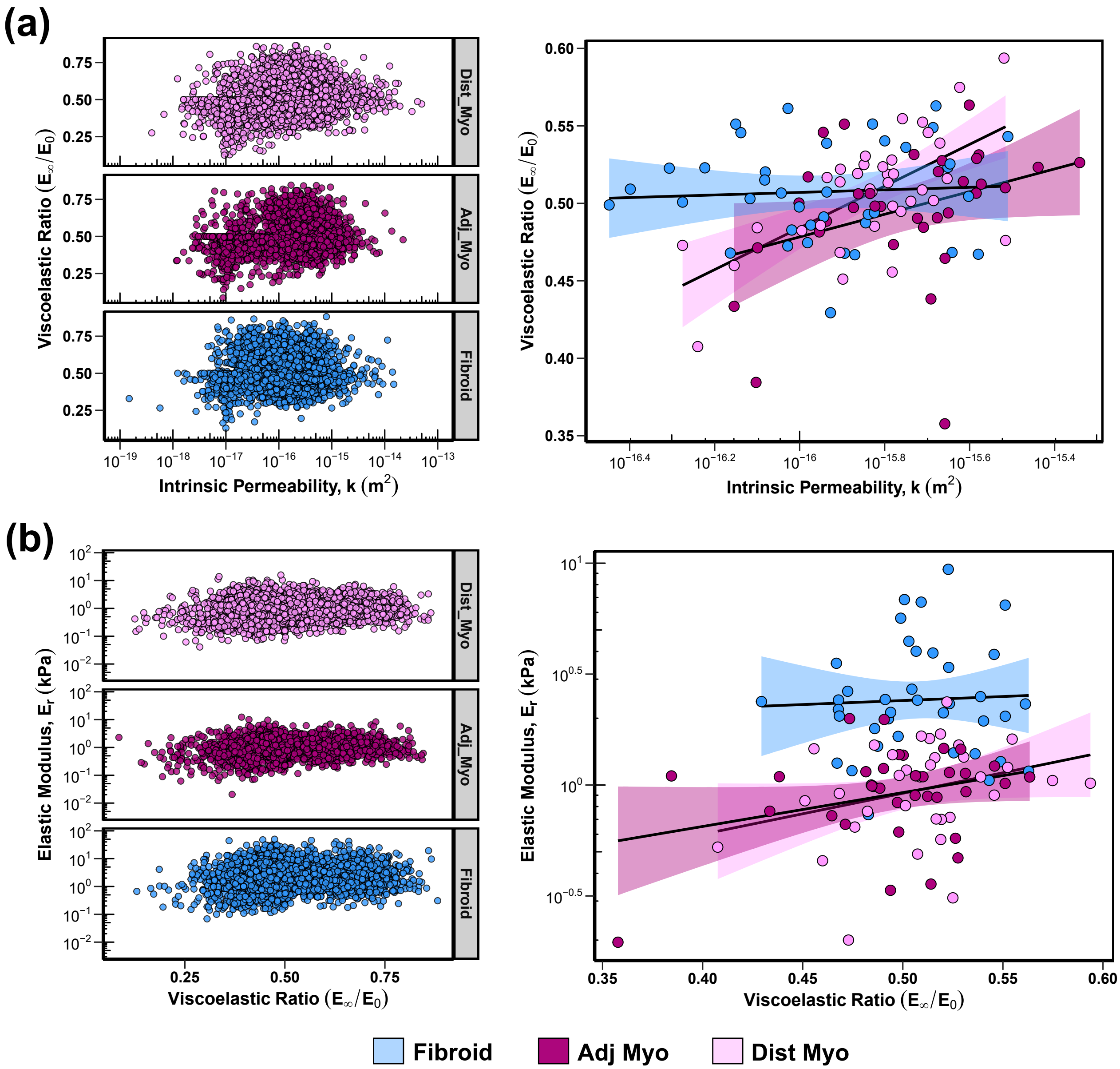


**Figure S10. Correlation of material properties for (a) viscoelastic ratio vs permeability and (b) elastic modulus vs permeability.** Data are shown for all technical replicates (left) and median values per sample (right).

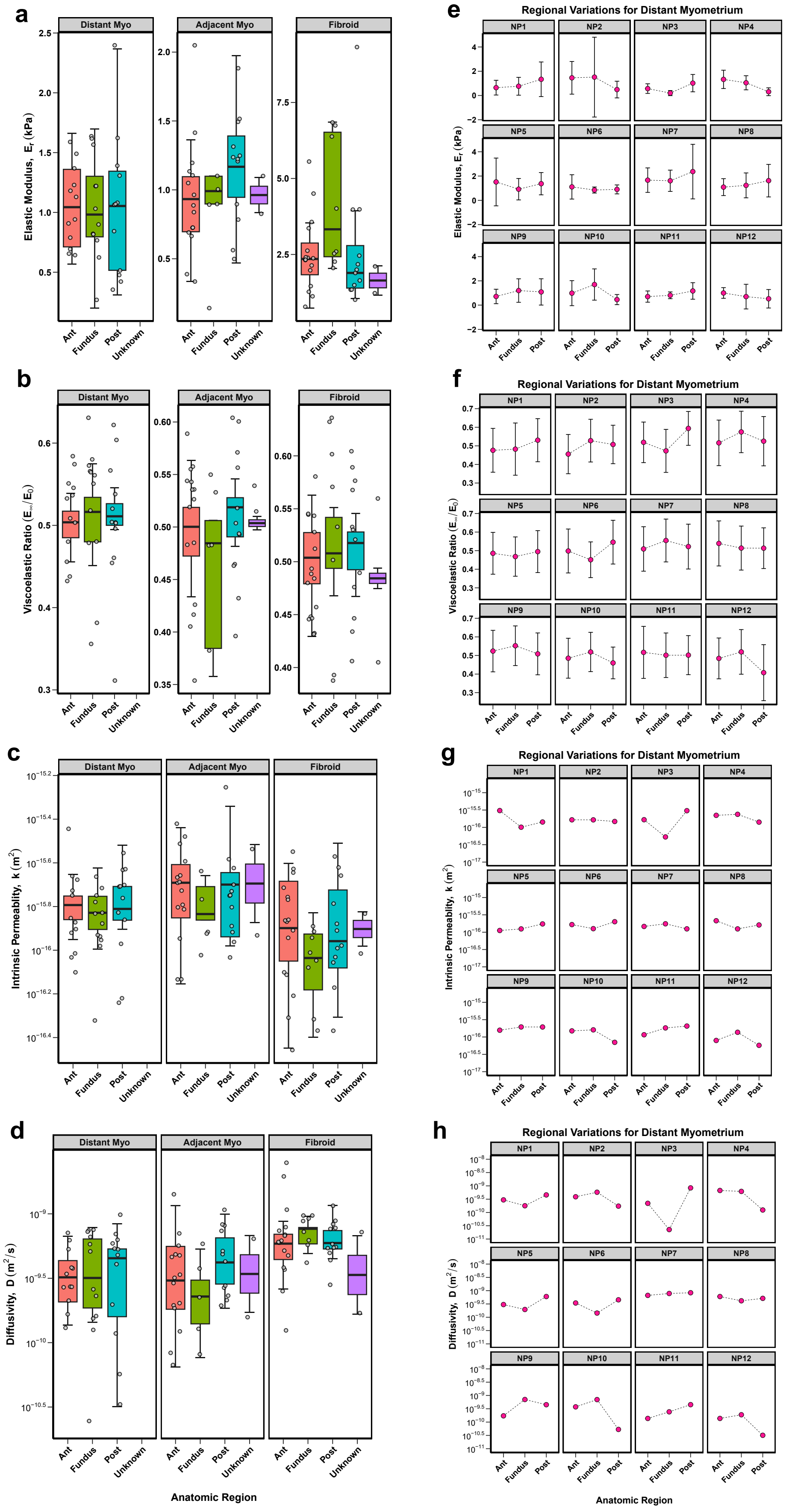


**Figure S11. Material properties by anatomic regions. (a-d)** Boxplots for distant myometrium, adjacent myometrium, and fibroid tissues. **(e-h)** Patient-matched data for distant myometrium are presented. Each symbol represents the mean value of all indentation points measured for an individual sample.

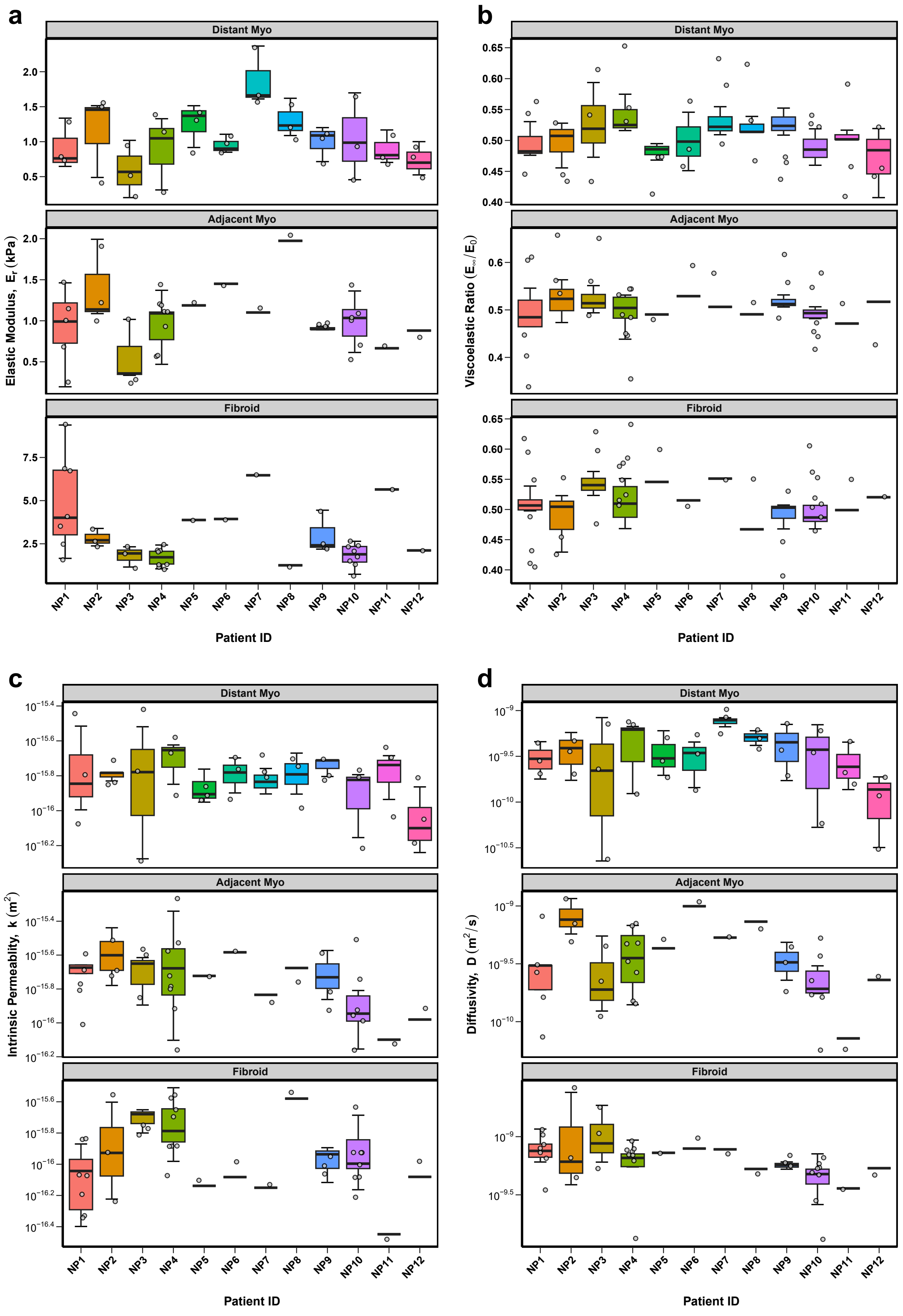


**Figure S12. Material properties plotted by patient for distant myometrium, adjacent myometrium, and fibroid tissues.** Data are shown for: (a) elastic modulus ($E_r$), (b) viscoelastic ratio ($E_{inf}/E_0$), (c) permeability ($k$), and (d) diffusivity ($D$). Each symbol represents the mean value of all indentation points measured for an individual sample.

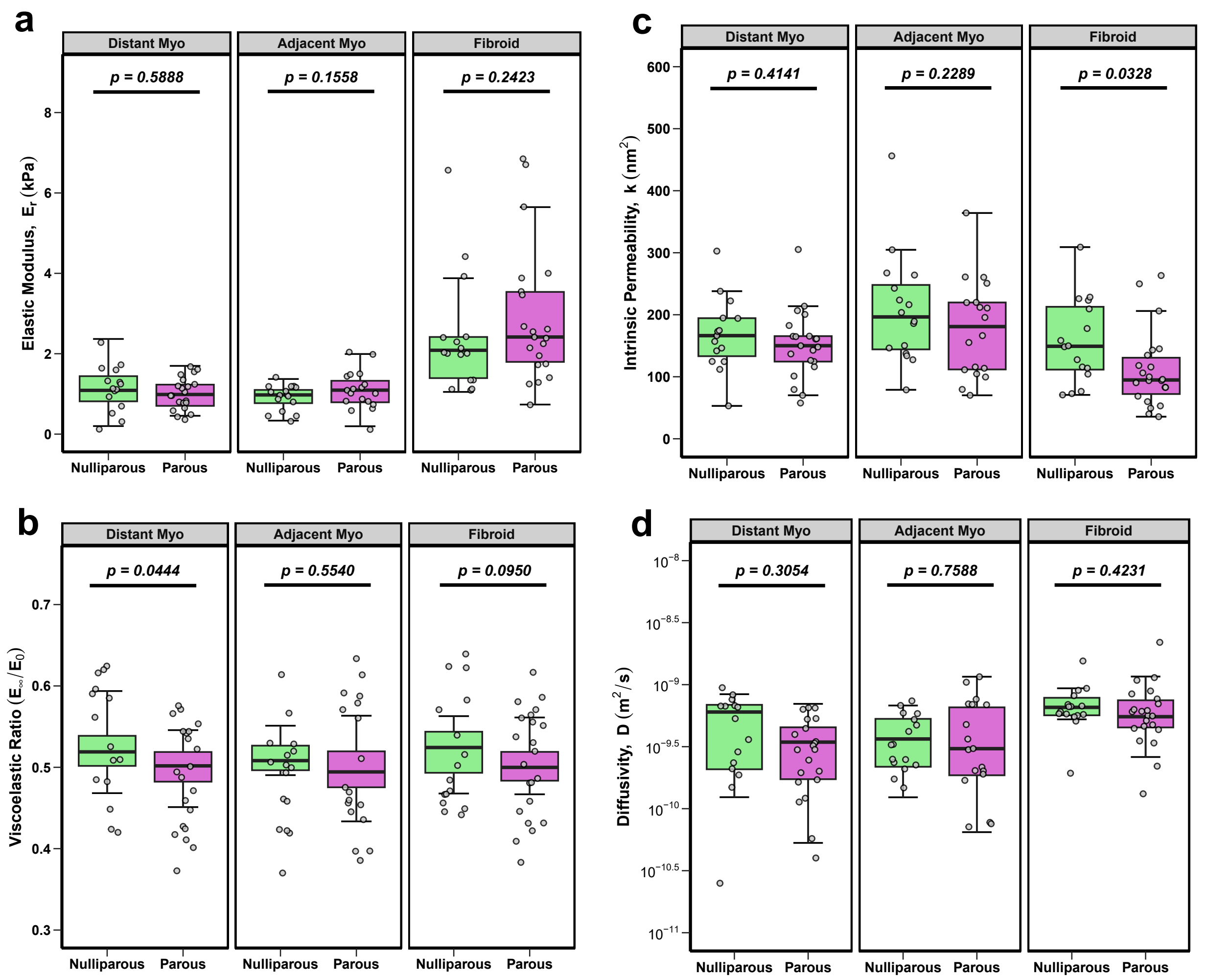


**Figure S13. Effect of parity on the material properties on distant myometrium, adjacent myometrium, and fibroid tissues.** Data are shown for: (a) elastic modulus ($E_r$), (b) viscoelastic ratio ($E_{inf}/E_0$), (c) permeability ($k$), and (d) diffusivity ($D$). Each symbol represents the mean value of all indentation points measured for an individual sample. Statistical analysis was performed using a linear mixed model, with sample ID included as a random effect; p-values are noted for each comparison.

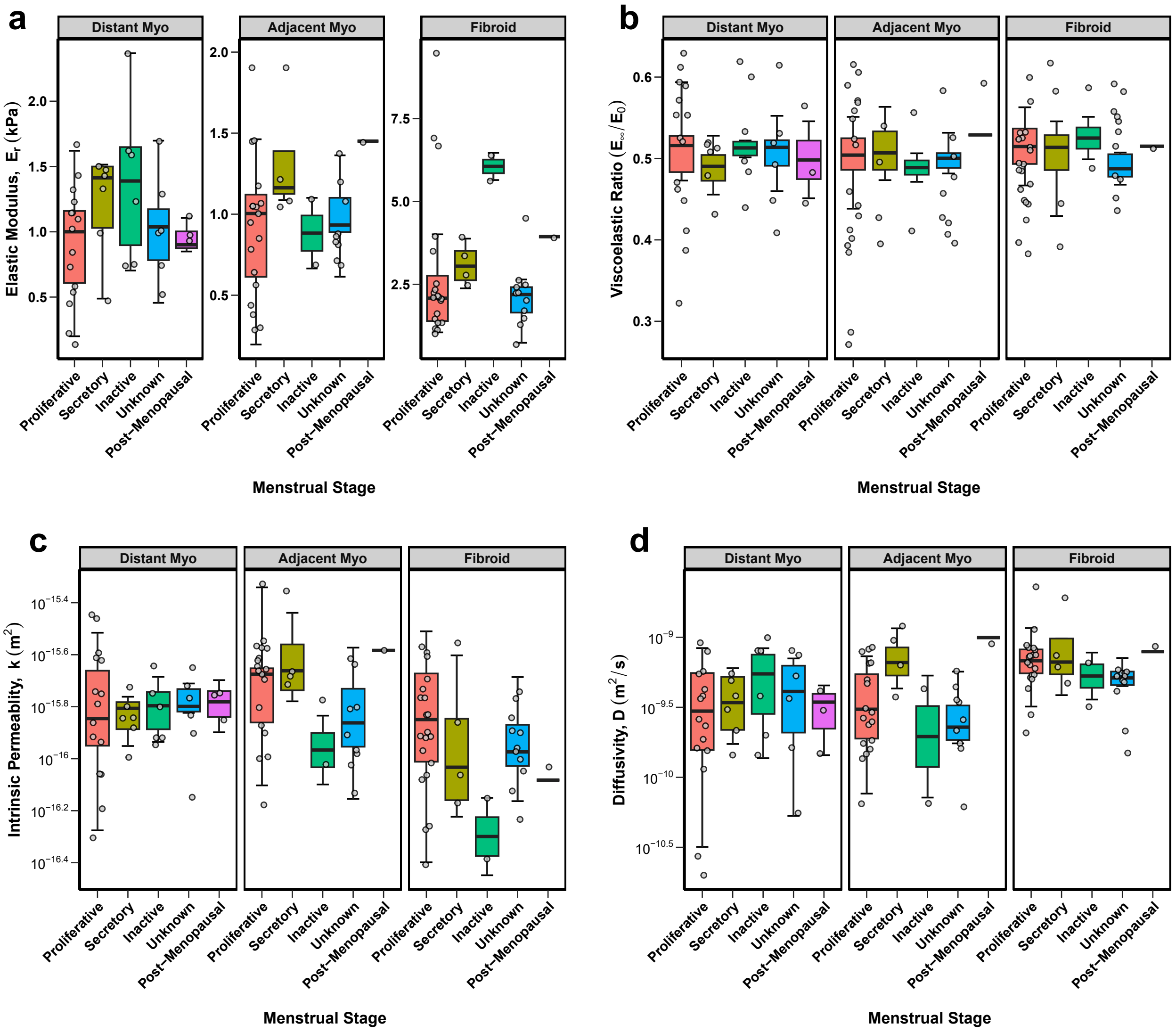


**Figure S14. Material properties by menstrual cycle phase for distant myometrium, adjacent myometrium, and fibroid tissues.** Data are shown for: (a) elastic modulus ($E_r$), (b) viscoelastic ratio ($E_{inf}/E_0$), (c) permeability ($k$), and (d) diffusivity ($D$). Each symbol represents the mean value of all indentation points measured for an individual sample.

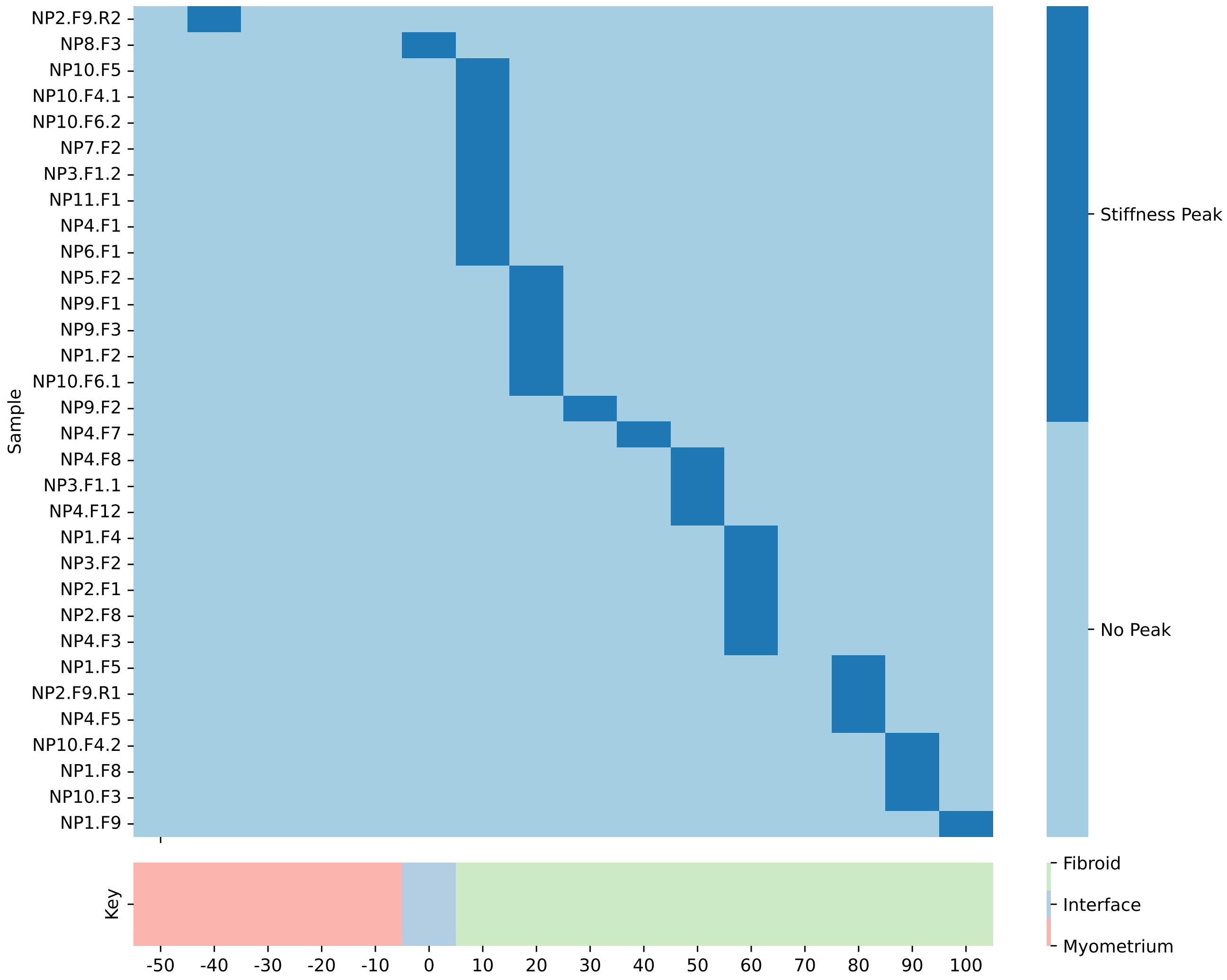


**Figure S15.** **Relative location of maximum stiffness (i.e. elastic modulus, $E_r$) on an individual sample basis.** Each row represents a single sample's testing region with paired fibroid (*green*) and myometrium (*red*) tissue. Only a single row of data at the central position of the testing region is included in this analysis. The x-axis is shown as a percentage of fibroid length. The interface position is marked at 0. Dark blue shaded squares indicate the positions of maximum stiffness. The distance between each discrete column of data is 200 $\mu$m.

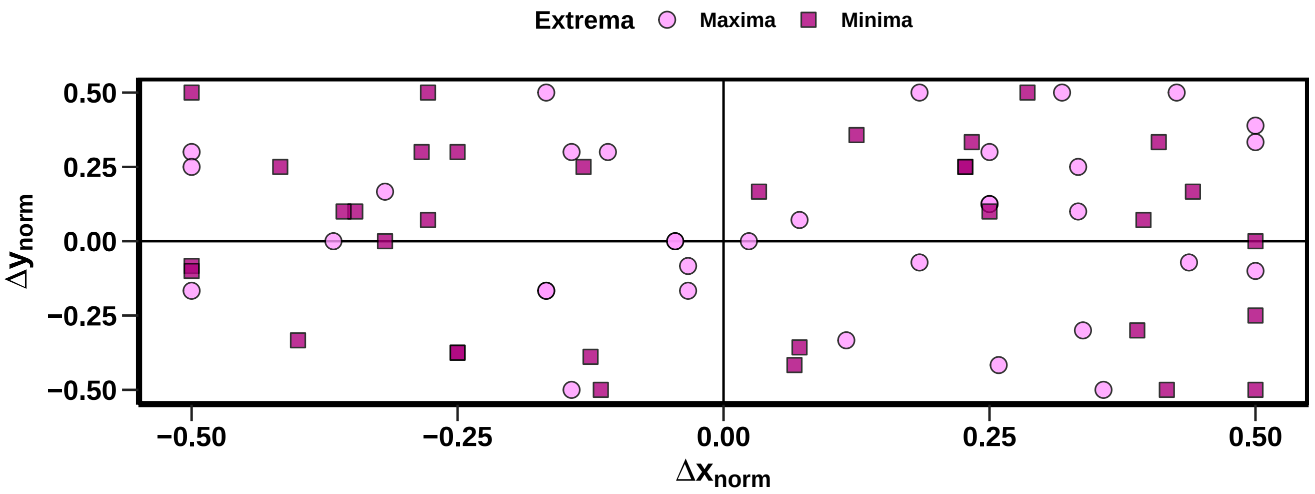


**Figure S16.** **Location of fibroid extrema relative to the fibroid's center (0, 0).** Each symbol represents the maxima (*circle*) or minima (*square*) elastic modulus ($E_r$) value of fibroid tissue. The positions ($\Delta y_{norm}$, $\Delta x_{norm}$) of extrema are normalized relative to the fibroid's length ($L_F$) and testing region width ($Y$).

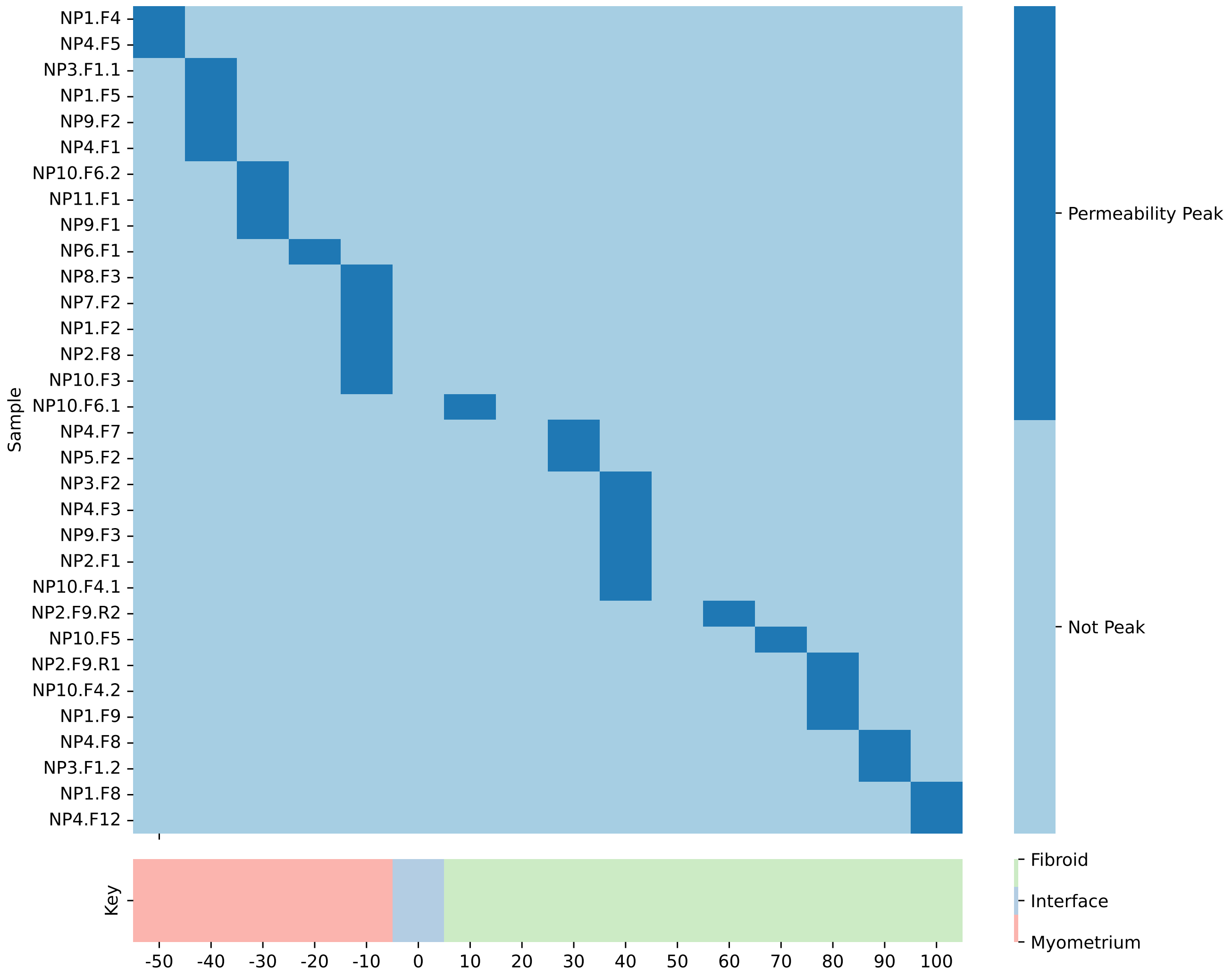


**Figure S17. Relative location of maximum permeability on an individual sample basis.** Each row represents a single sample's testing region with paired fibroid (*green*) and myometrium (*red*) tissue. Only a single row of data at the central position of the testing region is included in this analysis. The interface position is marked at X0. The X-axis is a percentage of fibroid length. Dark blue shaded squares indicate the positions of maximum stiffness. The distance between each discrete column of data is 200 $\mu$m.

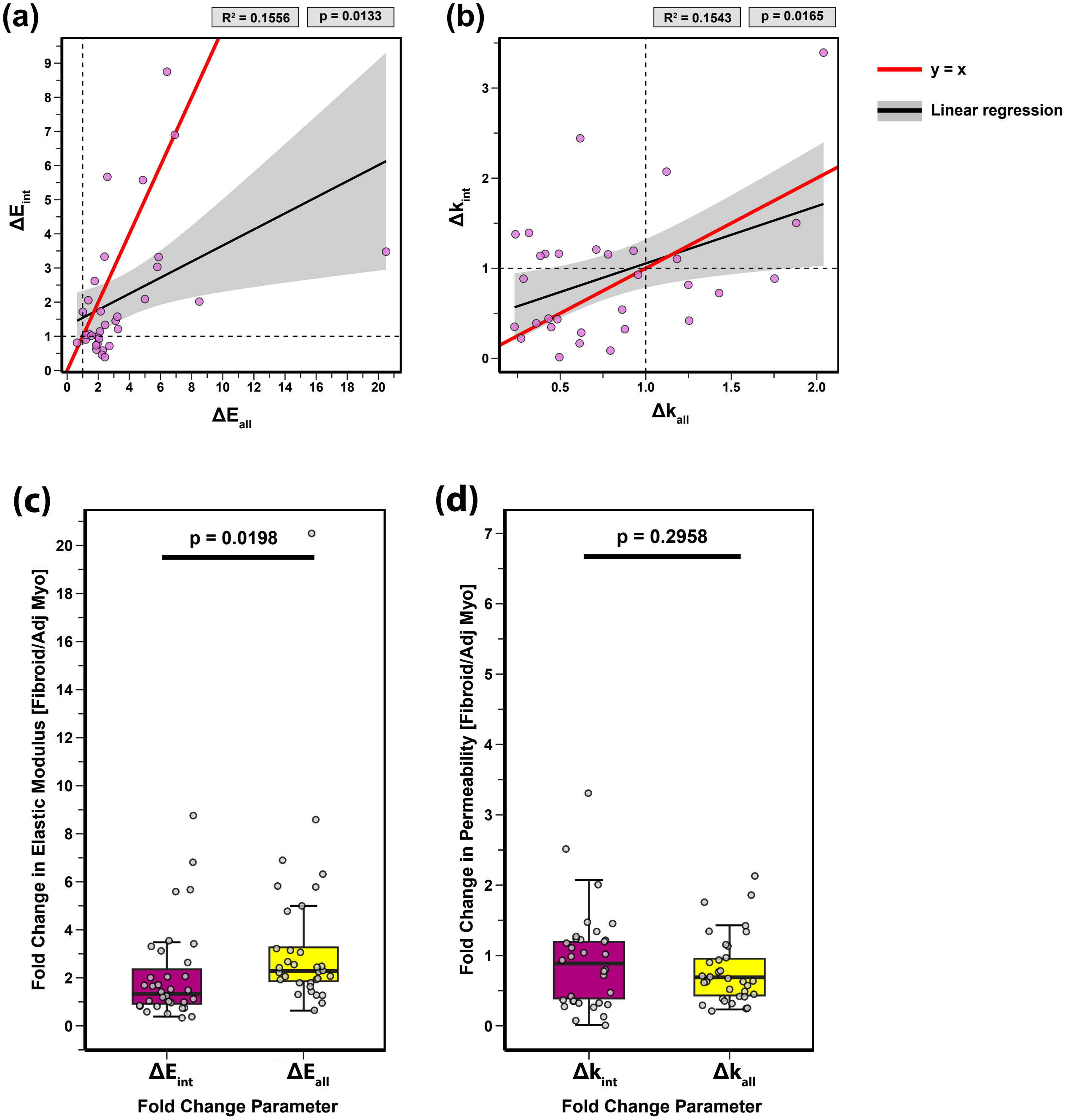


**Figure S18. Fold change parameters compared for (a, c) elastic modulus and (b, d) permeability**. Data are presented as x–y scatter plots (a, b) and box plots (c, d). $p$ and adjusted $R^2$ values are noted for each comparison. A solid red line marks a perfectly linear relationship ($y = x$). Linear regression lines are black, and standard deviations are shaded in grey. Each symbol represents a distinct sample for which the data of multiple indentation points have been averaged. Statistical analysis was performed using a linear mixed model, with patient ID included as a random effect; p-values are noted for each comparison.

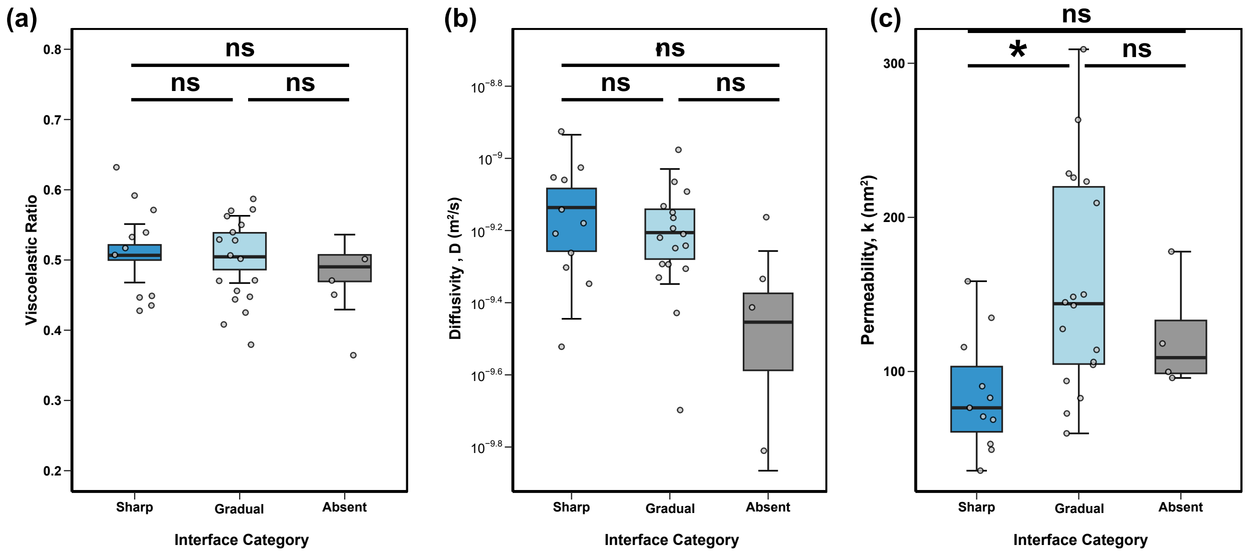


**Figure S19. Material parameters by elastic modulus interface categories.** Data is shown for (a) viscoelastic ratio, (b) diffusivity, and (c) permeability. Each symbol represents a distinct sample for which the data of multiple indentation points have been averaged.

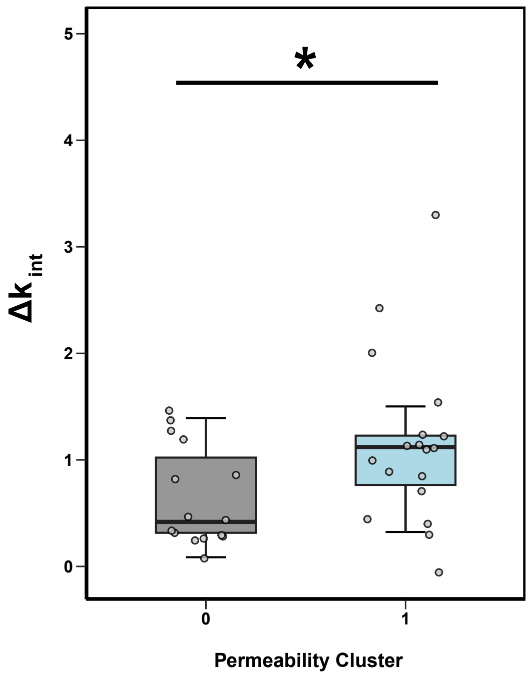


**Figure S20. Relative change in intrinsic permeability ($k_{int}$) calculated between fibroid and adjacent myometrium samples at the interface region versus the k-means cluster category.** Each symbol represents a distinct sample. Statistical analysis was performed using a linear mixed model, with sample ID included as a random effect; significance is denoted as follows: $^{*}$ $p < 0.05$, $^{**}$ $p < 0.01$.

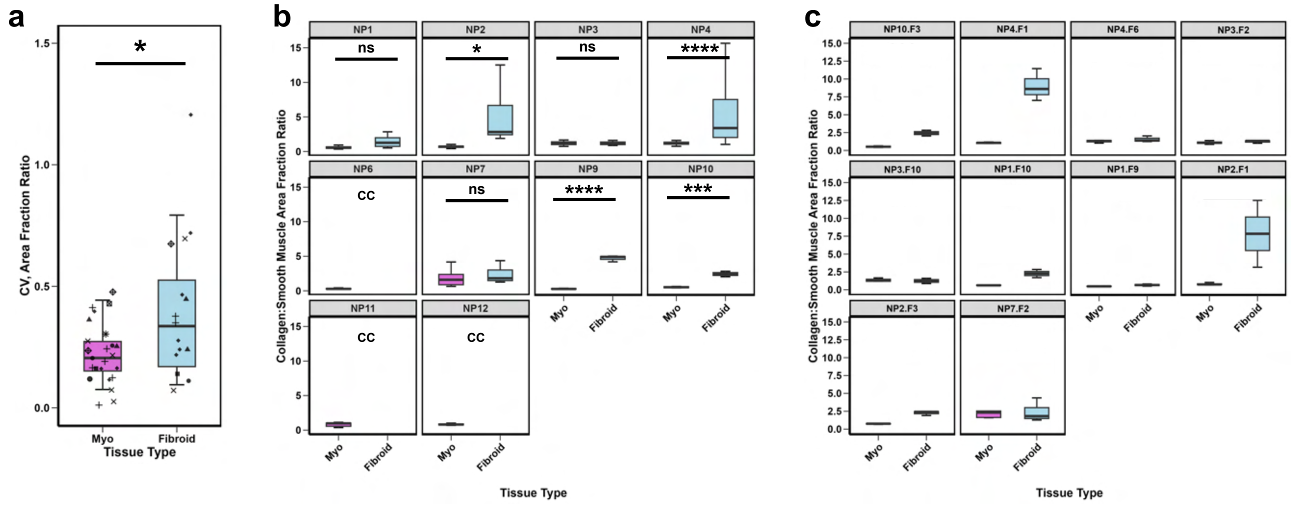


**Figure S21.** **Image quantification of collagen:smooth muscle area fraction ratios.** **(a)** Coefficient of variance (CV) by tissue type. Each distinct symbol corresponds to a distinct patient. Statistical analysis was performed using a linear mixed model, with sample ID included as a random effect; significance is denoted as follows: $^{*}$ $p < 0.05$. **(b)** Collagen:smooth muscle area fraction plotted by patient. T-tests with a Bonferroni corrections were used to compute statistical significance for each comparison. Significance is denoted as follows: $^{ns}$ $p > 0.05$, $^{*}$ $p < 0.05$, $^{**}$ $p < 0.01$, $^{***}$ $p < 0.001$, $^{****}$ $p < 0.0001$. Comparisons marked with CC indicate that data could not be computed where one group or more had less than two replicates. **(c)** Collagen:smooth muscle area fraction plotted on an individual fibroid sample basis.

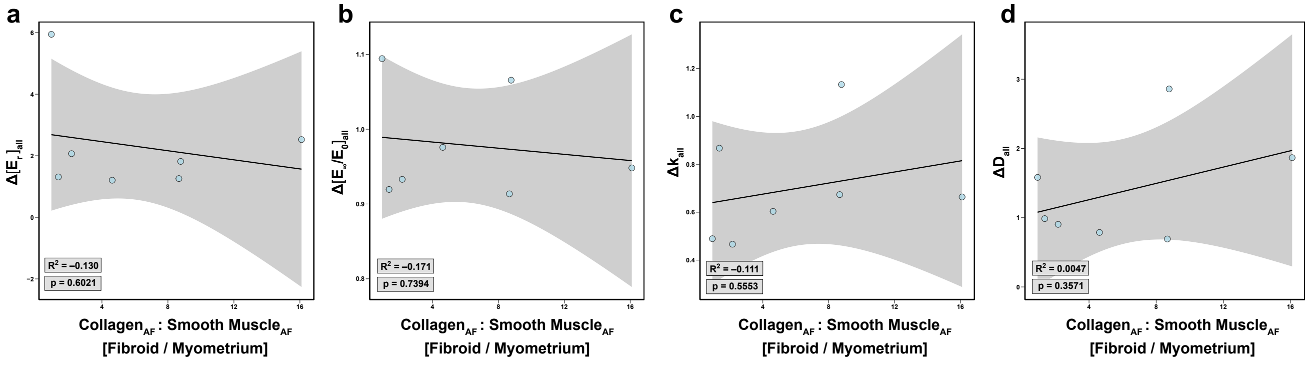


**Figure S22.** **Correlation of image analysis and material property data.** The relative change in the collagen:smooth muscle area fraction (AF) ratio between fibroid and myometrium tissues is plotted on the x axis. The relative change in material properties between fibroid and adjacent myometrium tissues is plotted on the y axis. Each distinct plot depicts a different material property: (a) $\Delta[E_r]_{all}$, (b) $\Delta[E_\infty/E_0]_{all}$, (c) $\Delta k_{all}$, (d) $\Delta D_{all}$. p-values and adjusted $R^2$ values from linear regression analysis are noted in the bottom left corner of each plot.

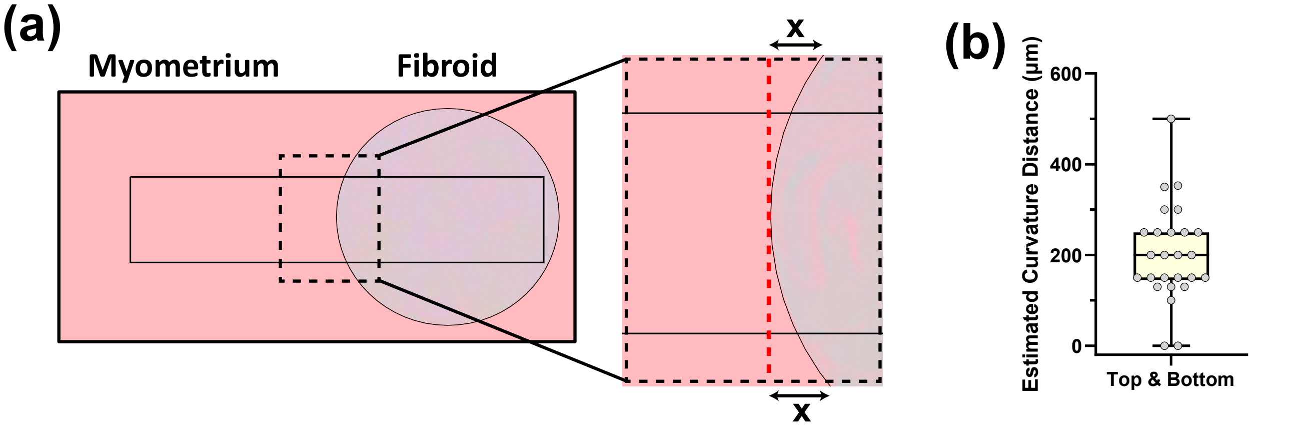


**Figure S23. Quantification of fibroid curvature within the rectangular testing region.** (**a**) Schematic of microindentation testing protocol for spatially mapping the fibroid-myometrium interface, zooming in on the interface region. The distance (x) between the leading edge of the fibroid (dashed red line) and the top-most and bottom-most edges of the fibroid within the testing region was quantified from dissecting microscope images for a subset of samples. **(b)** Boxplot of the estimated curvature distance (x), pooling data from the top and bottom edge measurements.

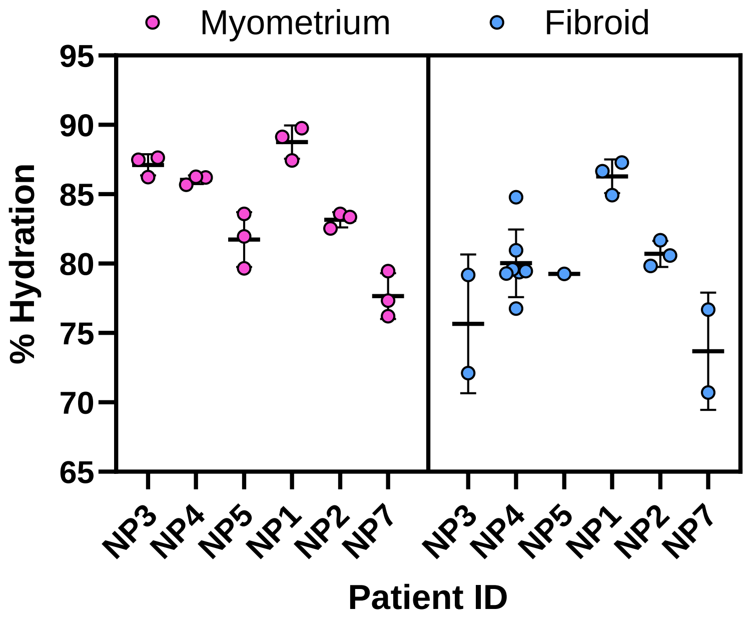


**Figure S24. Hydration (%) of myometrial and fibroid tissues by patient.** Each dot represents a single measurement taken for a single tissue sample.

**Table S3.** Summary of SHG imaging parameters for a subset of samples.

| Sample ID | Anatomic Region | No. Z-Stacks | Z-Stack Distance |
|---|---|---|---|
| NP1.F3–4 | Fundus | 3 | 10 $\mu$m |
| NP1.F5 | Fundus | 3 | 10 $\mu$m |
| NP1.F8 | Posterior | 5 | 15 $\mu$m |
| NP1.F9 | Posterior | 3 | 10 $\mu$m |
| NP1.Myometrium (Distant) | Anterior | 3 | 10 $\mu$m |

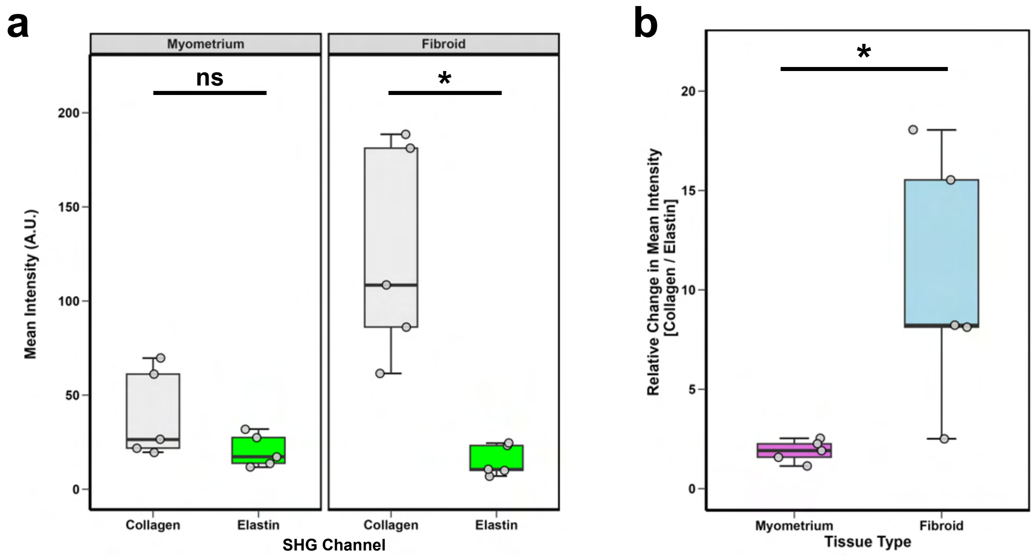


**Figure S25. Additional quantitative measures of SHG image analysis.** (a) Mean intensity of collagen (*860 nm*) and elastin (*1040 nm*) channels for myometrium and fibroid tissues, displaying arbitrary units (A.U.). Comparisons between SHG channels are shown. Each dot represents the mean value of technical replicates for a given sample. (b) Mean intensity ratio of collagen to elastin for myometrium and fibroid tissues. For all comparisons, statistical analysis was performed with a Welch's t-test with a Bonferroni correction with significance marked as follows: $^{ns}p > 0.05$, $^{*}p < 0.05$).

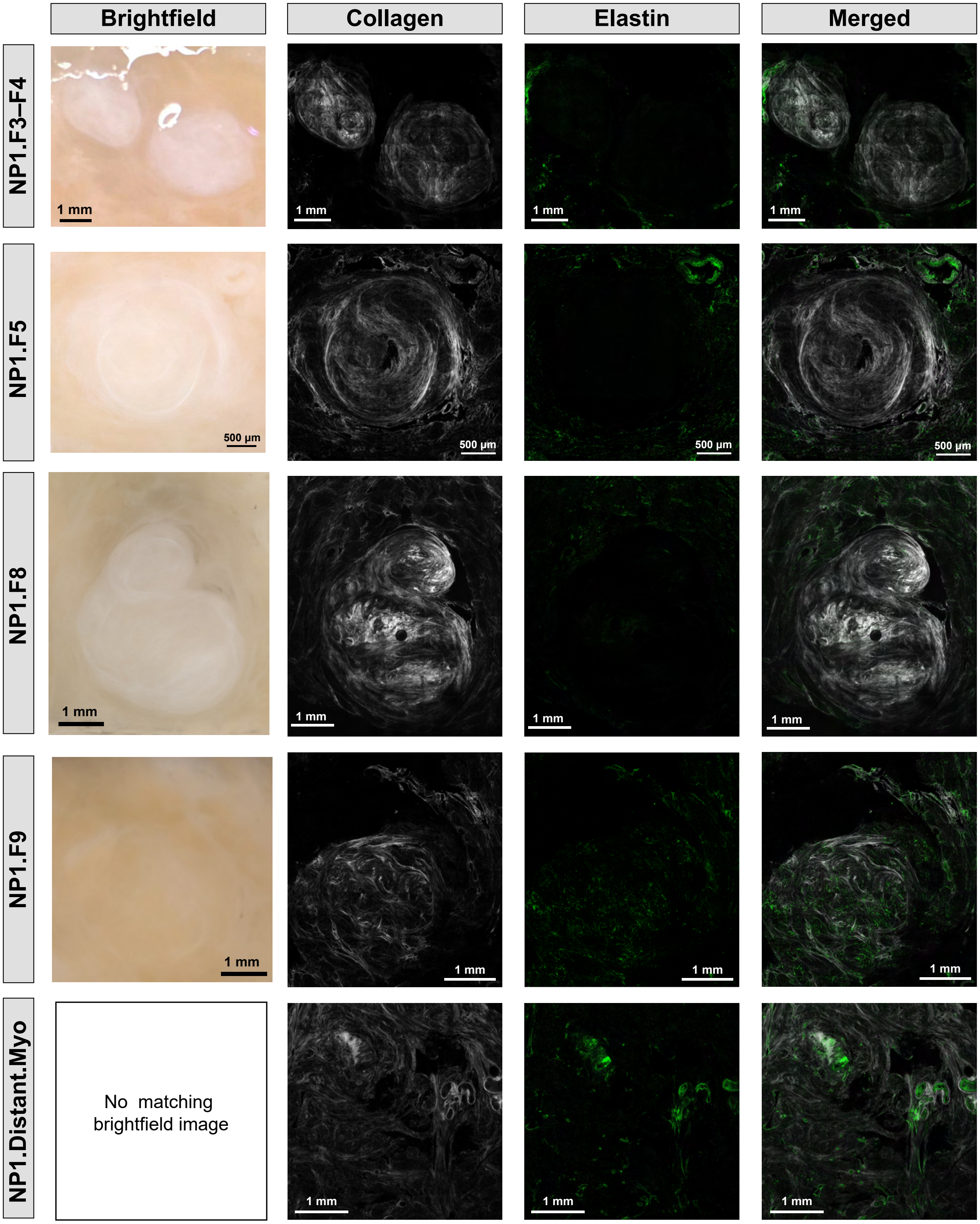


**Figure S26.** **Additional samples imaged with second harmonic generation (SHG) imaging.** Data represent four seedling fibroids and one sample of distant myometrium; each row represents the matched images for a given sample. SHG images (columns 2–4) and matching brightfield images (column 1) are shown for each sample. A single representative z-stack is shown.

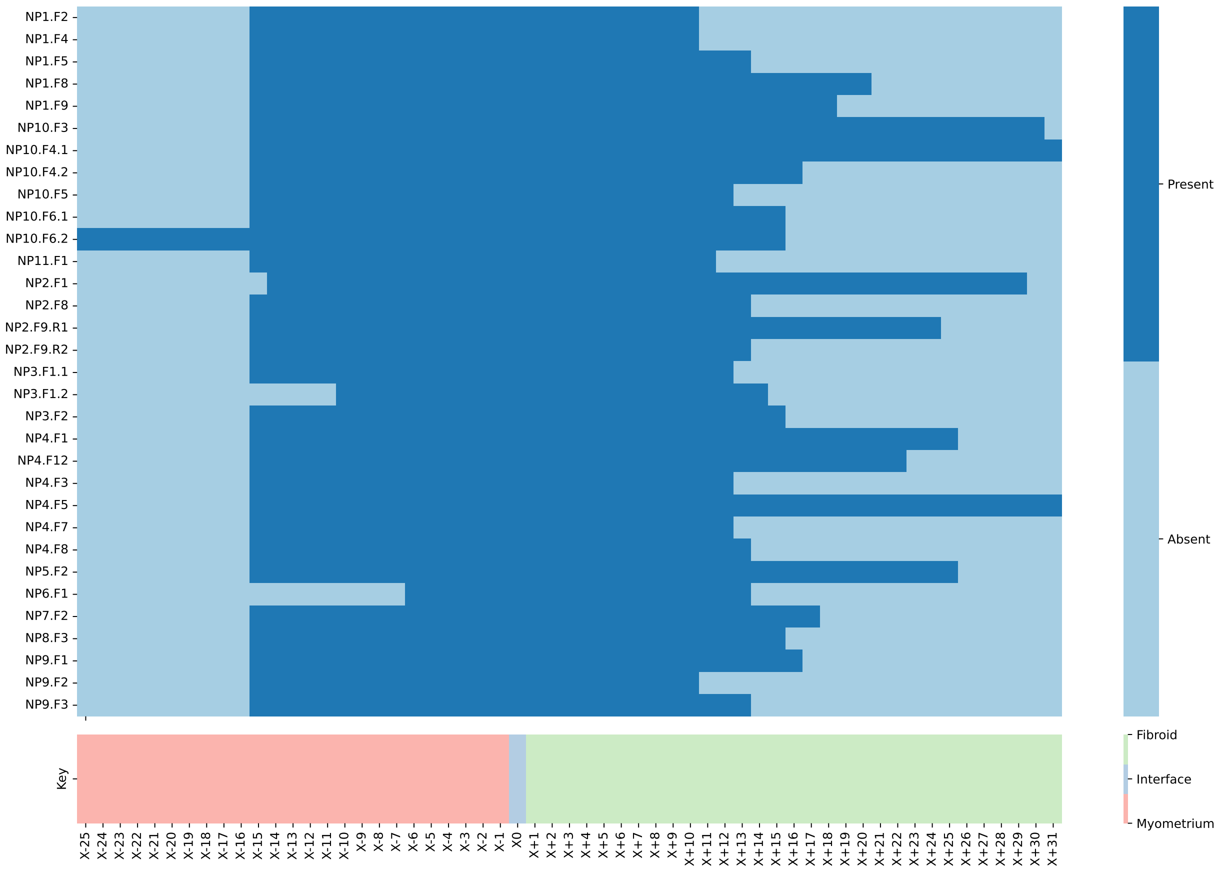


**Figure S27. Dimensionality of samples.** Heatmap showing presence and absence of samples. X- and X+ horizontal indices indicate myometrium and fibroid tissues, respectively. X0 marks the interface between the two regions. Dark blue boxes indicate tested horizontal indices; light blue boxes indicate an absence of measured values for a given index.

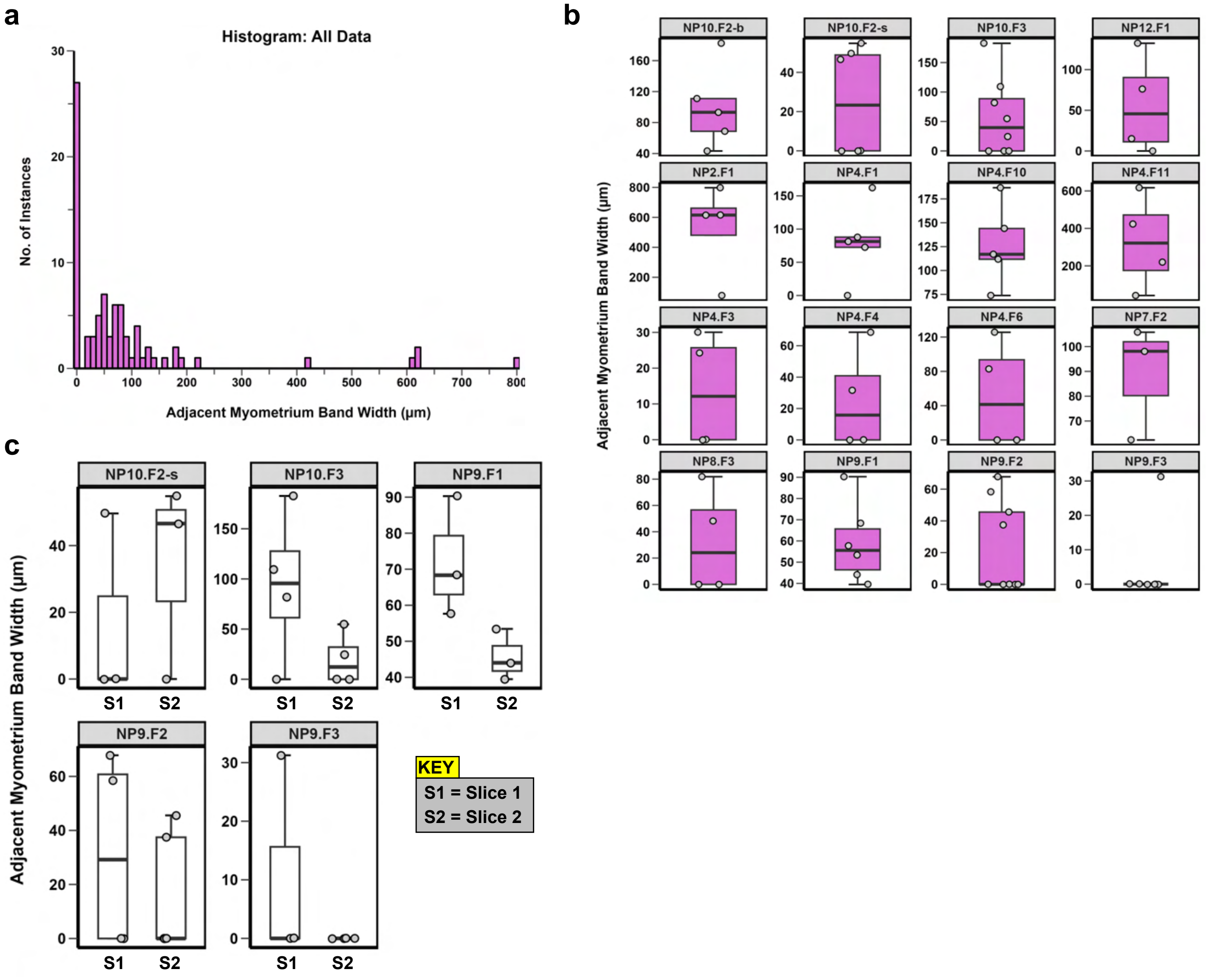


**Figure S28. Quantification of adjacent myometrial tissue band width.** **(a)** Histogram of all data. **(b)** Distribution of width measurements for each distinct fibroid. **(c)** Distribution of width measurements for identical samples measured for different depths of histological cuts (S1 = slice 1, S2 = slice 2). Distances between slices were variable and were not explicitly quantified.

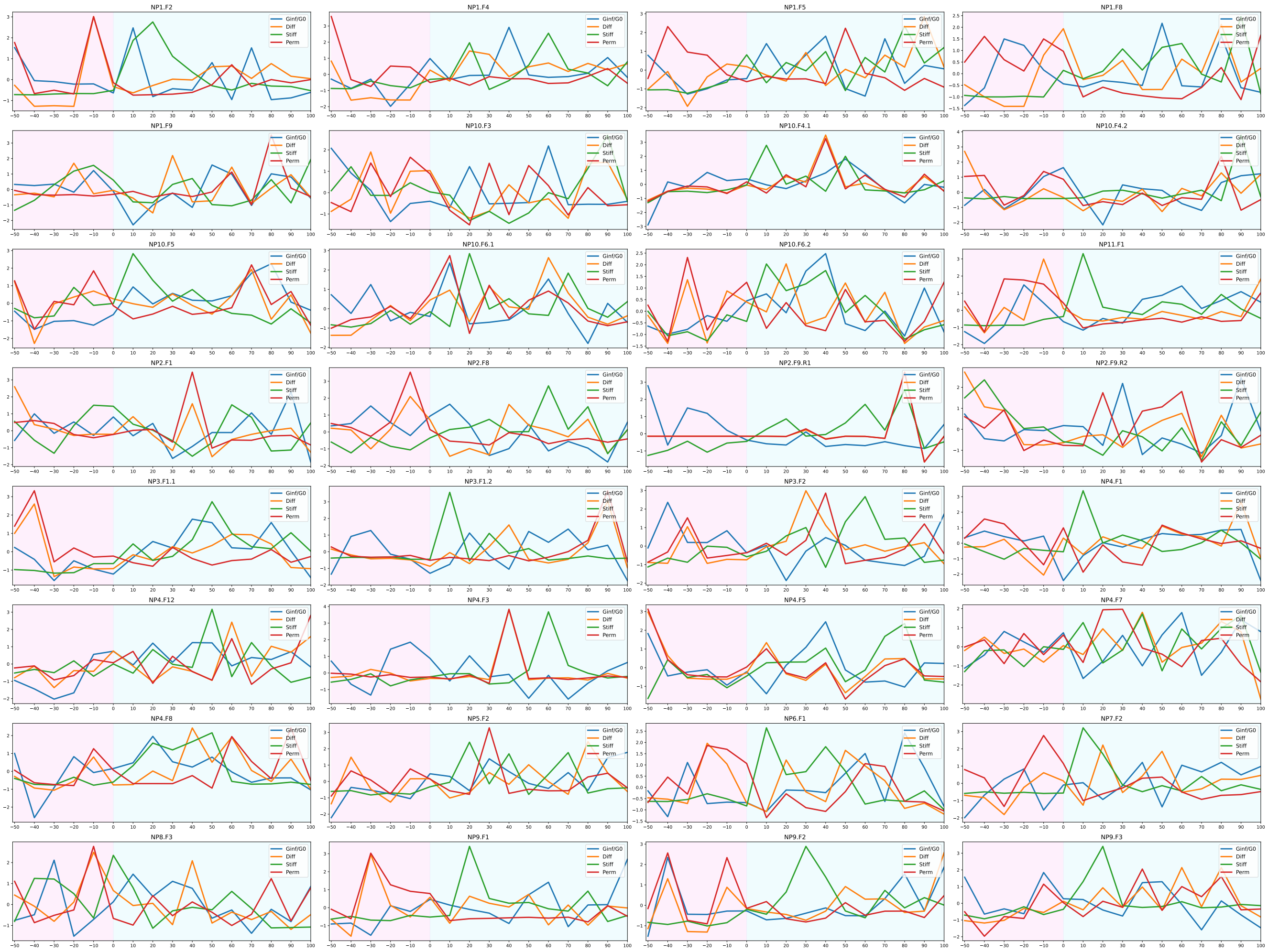


**Figure S29. Data inputs for k-means clustering.** Each plot represents a distinct specimen mapping across the adjacent myometrium (pink) and fibroid (blue) regions. Each line represents a distinct material parameter: viscoelastic ratio ($E_{inf}/E_0$), diffusivity ($D$), stiffness ($E_r$), and permeability ($k$). The data shown represents the central row of data. The x-axis is displayed as a percentage of fibroid length.

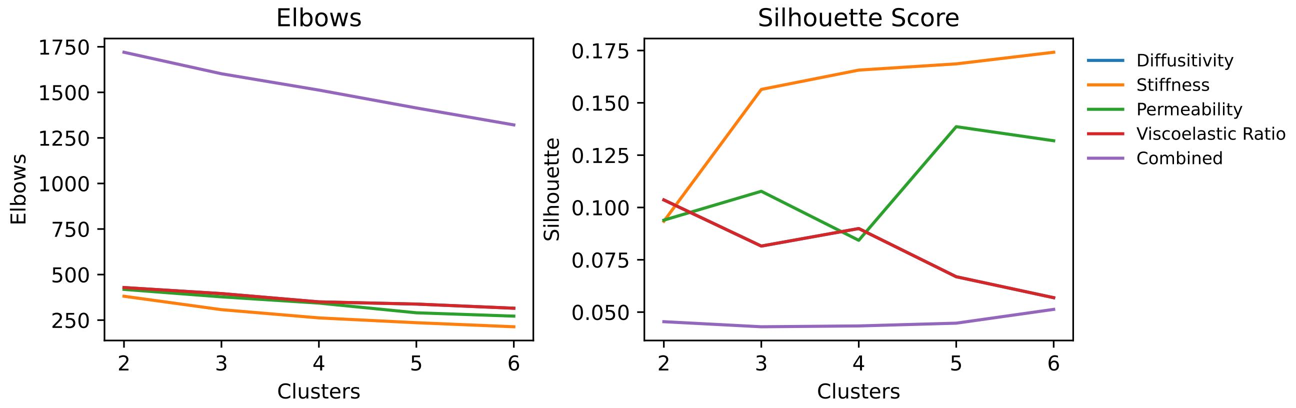


**Figure S30.** **Heuristic for determining number of clusters.** Elbow (left) and Silhouette (right) plots used to evaluate the quality of clusters. Each line represents a distinct material property (i.e., elastic modulus/stiffness, viscoelastic ratio, permeability, and diffusivity) or the combination of all (i.e., combined). Elbow plots show the within-cluster sum of squares against the number of clusters. Silhouette score plots demonstrate how similar a data point is to its own cluster compared to other nearby clusters.

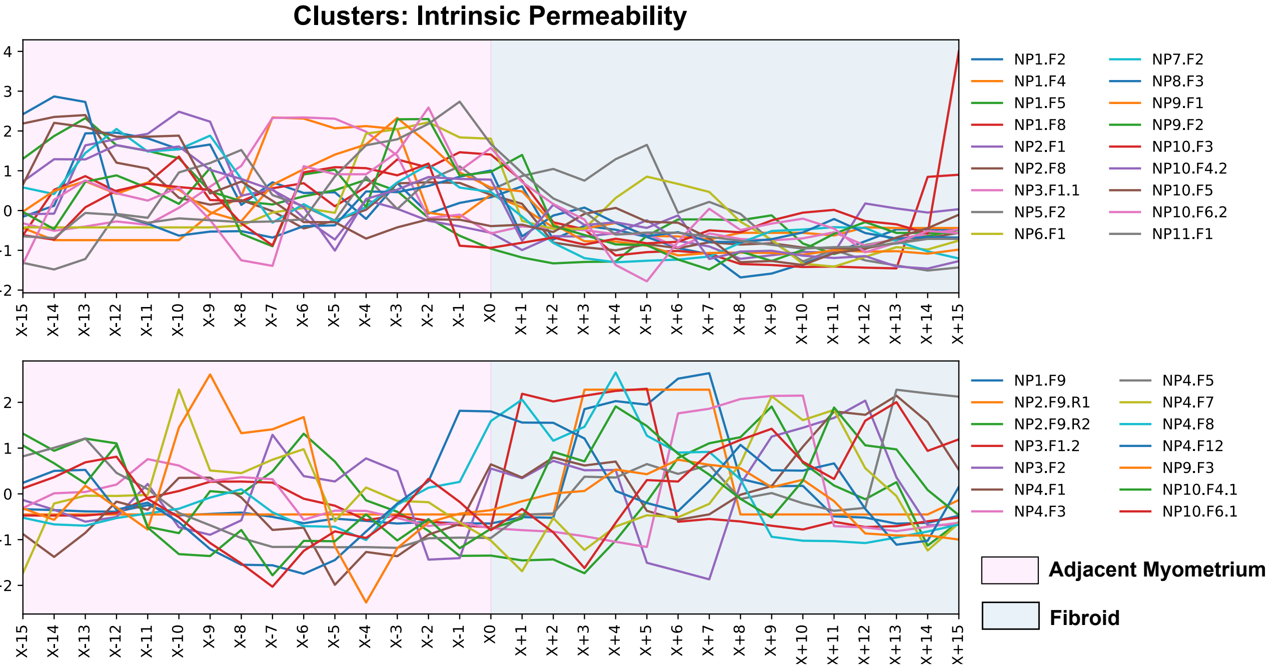


**Figure S31.** **K-means clusters by individual sample for intrinsic permeability (k) values.** Data are shown for cluster 0 (top) and cluster 1 (bottom). Adjacent myometrium regions are shaded in pink and include negative X values. Fibroid regions are shaded in blue and include positive X values. X0 marks the interface region. The y-axis displays arbitrary units.

| **Study ID** (*Fodera et al. 2025*) | **Other Study** | **Tissue Analyzed** | **Matching Patient ID** |
|---|---|---|---|
| NP1 | [61] Fang *et al.* 2025 | Myometrium | NP4 |
| NP2 | [45] Fodera *et al.* 2024<br>[61] Fang *et al.* 2025<br>[101] Therien *et al.* 2025 | Endometrium, Myometrium, Perimetrium<br>Myometrium<br>Uterine Fibroid & Myometrium | NP3<br>NP3<br>8 |
| NP3 | [61] Fang *et al.* 2025<br>[101] Therien *et al.* 2025 | Myometrium<br>Uterine Fibroid & Myometrium | NP2<br>3 |
| NP4 | [101] Therien *et al.* 2025 | Uterine Fibroid & Myometrium | 4 |
| NP5 | [101] Therien *et al.* 2025 | Uterine Fibroid & Myometrium | 7 |
| NP6 | [61] Fang *et al.* 2025<br>[101] Therien *et al.* 2025 | Myometrium<br>Uterine Fibroid & Myometrium | NP5<br>9 |
| NP7 | [101] Therien *et al.* 2025 | Uterine Fibroid & Myometrium | 10 |
| NP8 | [101] Therien *et al.* 2025 | Uterine Fibroid & Myometrium | 6 |
| NP9 | [101] Therien *et al.* 2025 | Uterine Fibroid & Myometrium | 1 |
| NP10 | [101] Therien *et al.* 2025<br>[100] McLean *et al.* 2020 | Uterine Fibroid & Myometrium<br>Unspecified/Mixed | 2<br>NP Patient 2 |
| NP12 | [45] Fodera *et al.* 2024<br>[61] Fang *et al.* 2025<br>[101] Therien *et al.* 2025 | Endometrium, Myometrium, Perimetrium<br>Myometrium<br>Uterine Fibroid & Myometrium | NP2<br>NP6<br>5 |

**Table S4.** Linker table of Patient IDs to other studies in which patient-matched uterine tissue was evaluated.
